# Rapid gene evolution in an ancient post-transcriptional and translational regulatory system compensates for meiotic X chromosomal inactivation

**DOI:** 10.1101/2021.08.25.457683

**Authors:** Shengqian Xia, Iuri M. Ventura, Andreas Blaha, Annamaria Sgromo, Shuaibo Han, Elisa Izaurralde, Manyuan Long

**Affiliations:** Department of Ecology & Evolution, The University of Chicago, Chicago, Illinois, USA; Department of Biochemistry, Max Planck Institute for Developmental Biology, Tübingen, Germany; Research Institute of Molecular Pathology (IMP), VBC, 1030, Vienna, Austria; Insitute of Molecular Biotechnology (IMBA), VBC, 1030, Vienna, Austria; CAPES Foundation, Ministry of Education of Brazil, Brasília, DF 70040-020, Brazil

**Author notes:** these authors contributed equally (Shengqian Xia, Iuri M. Ventura). these authors contributed equally (Andreas Blaha, Annamaria Sgromo).

**Keywords:** CAF40, Poseidon, Zeus, CCR4-NOT, MXCI, MSCI, New Gene Function

## Abstract

It is conventionally assumed that conserved pathways evolve slowly with little participation of gene evolution. Nevertheless, it has been recently observed that young genes can take over fundamental functions in essential biological processes, for example, development and reproduction. It is unclear how newly duplicated genes are integrated into ancestral networks and reshape the conserved pathways of important functions. Here, we investigated origination and function of two autosomal genes that evolved recently in *Drosophila*: *Poseidon* and *Zeus*, which were created by RNA-based duplications from the X-linked *CAF40*, a subunit of the conserved CCR4-NOT deadenylase complex involved in post-transcriptional and translational regulation. Knockdown and knockout assays show that the two genes quickly evolved critically important functions in viability and male fertility. Moreover, our transcriptome analysis demonstrates that the three genes have a broad and distinct effect in the expression of hundreds of genes, with almost half of the differentially expressed genes being perturbed exclusively by one paralog, but not the others. Co-immunoprecipitation and tethering assays show that the CAF40 paralog Poseidon maintains the ability to interact with the CCR4-NOT deadenylase complex and might act in post-transcriptional mRNA regulation. The rapid gene evolution in the ancient post-transcriptional and translational regulatory system may be driven by evolution of sex chromosomes to compensate for the meiotic X chromosomal inactivation (MXCI) in *Drosophila*.

## INTRODUCTION

The complex regulation of gene expression is essential for a proper cell function. Accurate spatial and temporal transcriptional activation and repression, as well as post-transcriptional regulation, is determined by intricate regulatory networks, often involving extensive signaling processes and the recruitment of multi-protein regulatory complexes. Thus, describing the components of such regulatory circuits, as well as understanding their role in the evolution of gene regulation can extend our comprehension of how organisms adapt and diversify over time (Erwin and Davidson, 2009; Halfon, 2017). Fundamental cellular functions, including basic regulatory processes common to distantly related organisms, are often assumed to be carried out by old conserved elements, whereas evolutionary young genes would be involved in more restrict, even dispensable, activities (Miklos and Rubin, 1996; Kondrashov, 2012). However, recent case studies have challenged this view, showing that young genes can be incorporated into ancestral regulatory networks, with major impact in the expression of numerous genes (Matsuno et al., 2009; Ding et al., 2010; Chen et al., 2012). The importance of such integration of new elements into fundamental cellular processes is illustrated by dramatic examples of young genes which were experimentally shown to have acquired indispensable roles in development or reproduction in a short evolutionary time, even when present in a single or a small clade of species (Chen et al., 2010; Saleem et al., 2012; Ross et al., 2013; Lee et al., 2018; VanKuren and Long, 2018; Kasinathan et al., 2020).

The incorporation of new genetic elements, in particular evolutionary young genes, into ancestral regulatory networks remains elusive and underexplored (Abrusán, 2013; Zhang et al., 2015). In particular, little is known about the molecular mechanisms involved in the evolution of regulatory networks driven by recently evolved gene duplicates. For instance, it is not clear to what extent the regulatory role of a young gene diverges from that of the parental copy, as well as what specific cellular processes and phenotypes they impact. In this context, the detailed comparison of old genes and their young duplicated paralogs can shed light on the mechanisms leading to the integration of new elements into preexisting cellular processes.

Extensive comparative genomic analyses have revealed an intriguing evolutionary pattern of gene traffic: there is a strong excess of parental genes on the X chromosome that produced autosomal duplicated genes with specific expression in the male germline (Betrán et al., 2002; Vibranovski et al., 2009a; Kaessmann et al., 2009). This pattern was confirmed for various organisms such as flies (Betrán et al., 2002; Dai et al., 2006; Bai et al., 2007; Zhang et al., 2010), mosquitos (Toups and Hahn, 2010), and mammals (Emerson et al., 2004; Carelli et al., 2016). The preferential fixation of male-biased duplicated copies into autosomes likely reflects the fact that, during spermatogenesis, the silencing of X chromosome in meiosis and later stages, as observed in mammals (Richler et al. 1992) and *Drosophila* (Vibranovski et al., 2009b; Mahadevaraju et al., 2021) provided a selective mechanism to drive the X to A gene traffic in new gene duplication (Vibranovski et al., 2009a; Jiang et al., 2017; Long and Emerson et al., 2017). Consequently, natural selection favors the fixation of autosomal duplicate copies that escape the X chromosome and compensate for the expression of its parental gene (Betrán et al., 2002; Vibranovski et al., 2009a, 2012; Casola and Betrán, 2017). In addition, other factors were proposed to interpret the gene traffic out of the X. It was considered that the testis was a rapidly evolving organ, prone to the accumulation of new elements, consistent with intense sexual selection (Harrison et al., 2015). It was also shown that during late spermatogenesis stages, the transcription of new gene copies is facilitated by the permissive chromatin state, which may facilitate the promiscuous transcription, insertion, and subsequent evolution of newly arisen genes (Kaessmann, 2010; Witt et al., 2019).

Consistent with the pattern described above, previous studies have reported that male reproductive tissues express specific versions of several housekeeping genes involved in basic cellular processes, such as the proteasomal (Zhong and Belote, 2007), transcriptional (Hiller, 2004), and translational machineries (Baker and Fuller, 2007). It was suggested that these duplicated copies may represent specialized versions of their parental genes, required to accomplish the intense and coordinated changes in gene expression observed during spermatogenesis (White-Cooper, 2010). It is not clear, however, why the duplicated, specific copies occur so frequently, or to what extent they diverge in function from their parental ones (Belote and Zhong, 2009).

In this study, we investigate evolutionary and functional impacts of *Poseidon* and *Zeus*, two autosomal young genes expressed in the testes, which independently retroposed from X-linked *CAF40*, and are only found in some *Drosophila* species (Zhang et al., 2010). *CAF40* is an ancient gene, broadly expressed in all fly tissues. The locus encodes a highly conserved protein in eukaryotes, with orthologs identified and experimentally studied from mammals (e.g. mouse and human) to insects (e.g. *Drosophila*) to fungi (e.g. yeast) (Collart and Panasenko, 2017). CAF40, also known as Rcd-1 or CNOT9, is a subunit of the highly conserved CCR4-NOT deadenylase complex, a multi-protein assembly involved in post-transcriptional and translational regulation of gene expression (Miller and Reese, 2012; Wahle and Winkler 2013; Buschauer et al., 2020). The complex catalyzes the removal of poly(A) tails in mRNAs, thus leading to their translational repression and degradation. It is also involved in the deadenylation and degradation of mRNA targets for proper spermatogenesis (Legrand and Hobbs, 2018). By integrating several regulatory processes, the complex is considered a key regulator of eukaryotic gene expression (Collart, 2016). Among the subunits of the CCR4-NOT complex, CAF40 acts as an important hub for the recruitment of the complex by mRNA-binding proteins (Sgromo et al., 2017, 2018; Keskeny et al., 2019). Moreover, it was shown to act independently of the complex, by interacting with transcription factors and altering their activation potential (Garapaty et al., 2008).

*Zeus,* retroposed 5 million years ago (MYA), was already shown to play an important role in male fertility in *Drosophila*, by binding and regulating the expression of a large set of target genes, many not shared with the parental *CAF40* (Chen et al., 2012). In order to uncover how newly duplicated genes are integrated into ancestral networks and reshape the conserved pathways of important functions, we first describe the divergence between *CAF40* and its two retroduplicated genes, *Poseidon* and *Zeus*, in gene sequence and expression patterns. Second, using RNAi-knockdown and CRISPR-Cas9-deletions, we further explore their phenotypic importance for viability and male fertility. Third, we demonstrate that both duplicates are able to repress a tethered mRNA reporter, and that Poseidon protein, but not Zeus, retained the ability to interact with the CCR4-NOT complex. Finally, our RNA- seq data demonstrate that the independent silencing of each paralog impacts the regulation of a distinct set of genes, likely due to diverse functions between regulatory processes in which the paralogs are acting in. Together, our data show that both young duplicates are integrated into *Drosophila* male germline regulatory pathways, interact with highly conserved regulatory mechanisms, and impact the gene expression network in different ways.

## RESULTS

### Rapid evolution of *Poseidon* and *Zeus* out of the extremely conserved *CAF40*

Comparative genomic analyses from public databases had previously identified *Zeus* as a diverged copy of *CAF40* (Zhang et al., 2010), which prompted its functional description as a duplicated gene (Quezada-Díaz et al., 2010; Chen et al., 2012). Curiously, our analyses using the *CAF40* sequence as query in sequence search against *Drosophila* genomes had also revealed the presence of another annotated paralogous gene, although it had not been studied until now (*CG2053* in *D. melanogaster*). We named this gene *Poseidon*, as a reference to Zeus’ brother in the Greek mythology (PSI-BLAST e-value = 7x10^-56^, coverage = 81% between *CAF40* and *Poseidon* in *D. melanogaster*).

Our search revealed that the intact Open Reading Frames of *Poseidon* are present in the third chromosome of 18 *Drosophila* species (Fig. 1A), all from the subgenus *Sophophora*. Reciprocal BLAST searches using the *Poseidon* sequence as query did not find any other significant match in eukaryotes except for *CAF40* and *Zeus* orthologs. The phylogenetic distribution suggests that *Poseidon* is a relatively young gene that appeared 36 MYA, before *D. willistoni* diverged from other *Sophophora* species(Russo et al. 1995; Markow and O’Grady, 2007; Clark et al., 2007). *Zeus* originated after the split of the most recent common ancestor of *D. melanogaster* and *D. yakuba* 3-6 MYA (Quezada-Díaz et al., 2010; Chen et al., 2012; Russo et al. 1995; Markow and O’Grady, 2007; Clark et al., 2007). Despite the recent origination of these two new genes from different *CAF40* ancestries, these genes show a high level of divergence in their protein sequences (Fig. S1A).

**Figure 1.**
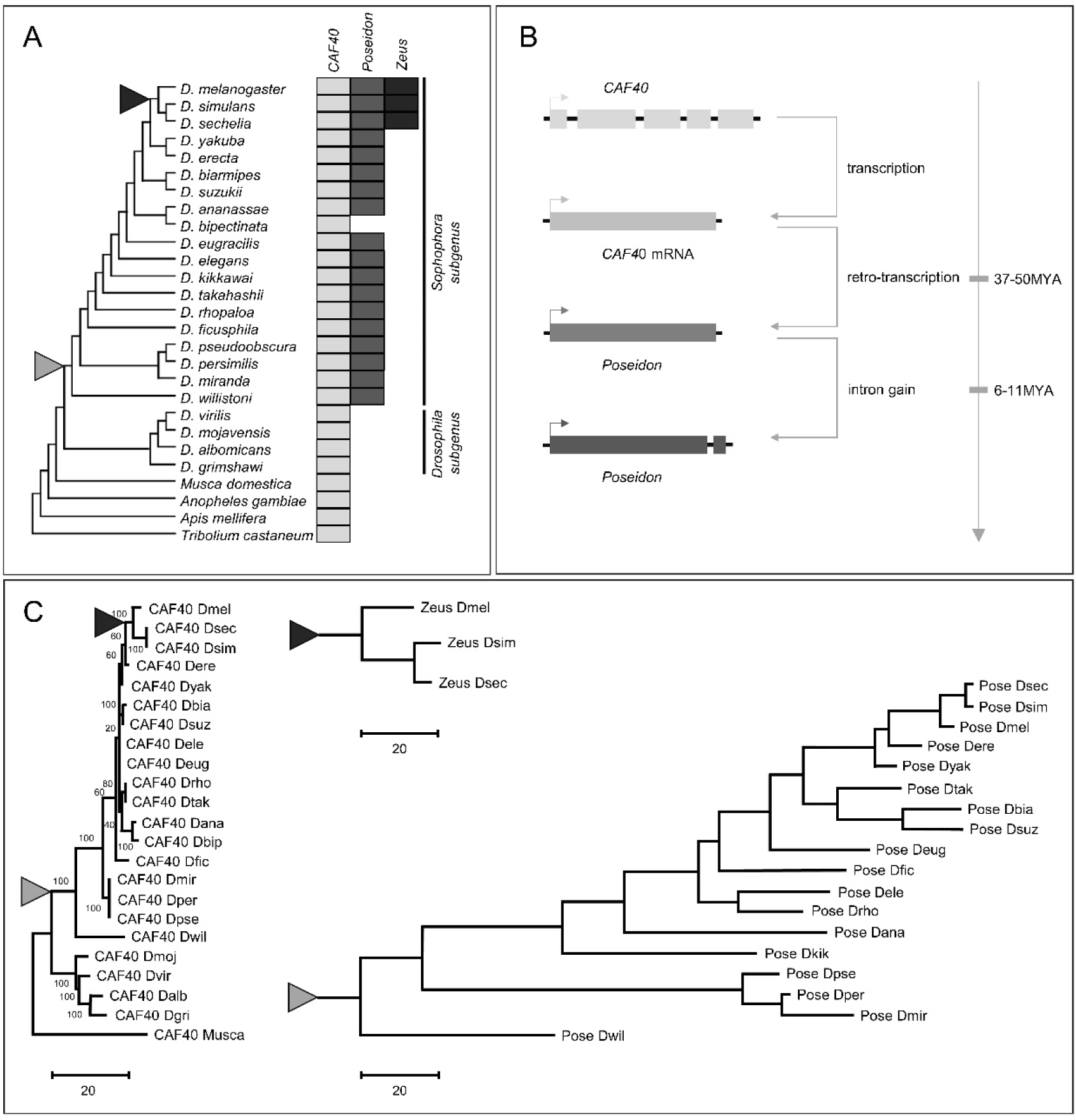
Origination of *Poseidon* and *Zeus* from *CAF40* in *Drosophila.* A.) Distribution of the three paralogs among *Drosophila* species, with other insects as outgroups. Filled boxes represent the presence of each gene. B.) Scheme depicting *Poseidon* origination process through retroduplication, and insertion into an autosome. C.) Phylogenetic relationship in *CAF40, Poseidon* and *Zeus* reconstructed respectively through the Maximum Parsimonious method using protein sequences. *Dmel – Drosophila melanogaster, Dsec – Drosophila sechellia, Dsim – Drosophila simulans, Dere – Drosophila erecta, Dyak – Drosophila yakuba, Dbia – Drosophila biarmipes, Dsuz – Drosophila suzukii, Dele – Drosophila elegans, Deug – Drosophila eugracilis, Drho – Drosophila rhopaloa, Dtak – Drosophila takahashii, Dana – Drosophila ananassae, Dbip – Drosophila bipectinate, Dfic – Drosophila ficusphila, Dmir – Drosophila miranda, Dkik – Drosophila kikkawai, Dper – Drosophila persimilis, Dpse – Drosophila pseudoobscura, Dwil – Drosophila willistoni, Dmoj – Drosophila mojavensis, Dvir – Drosophila virilus, Dalb – Drosophila albomicans, Dgri – Drosophila grimshawi, Musca – Musca domestica (Housefly).* The black and gray triangles represent the origination branch of *Zeus* and *Poseidon*, respectively. The scale for branch lengths is number of substitutions per site.

The presence of introns is useful for determining the mechanisms of origination of new genes (Long et al., 2013). *CAF40* orthologs have between 4 and 6 introns in *Drosophila* species. *Poseidon*, however, has no introns in all detected species except one small intron in its 3′end in the *D. melanogaster* subspecies group, which is unrelated in sequence or position to any intron found in *CAF40*. The lack of ancestral introns in the duplicated gene suggests that *Poseidon* initially originated through an X-to-autosome retroposition event, with the insertion of the duplicate in the third chromosome. Subsequently in the most recent common ancestor of the *D. melanogaster* subgroup species that diverged 6∼11 MYA, *Poseidon* gained a new intron (Fig. 1B).

Phylogenetic analyses of the three genes using the Maximum Parsimonious method in the MEGA platform (Kumar et al., 2018) suggest that *Poseidon* and *Zeus* originated through two independent RNA-based duplications from *CAF40* in different branches of the *Drosophila* phylogeny (Fig. 1C). The phylogenetic position of *Zeus* is consistent with its retroduplication after the split of the most recent common ancestor of *D. melanogaster* and *D. yakuba* (3-6MYA), as previously described (Quezada-Díaz et al., 2010; Chen et al., 2012). The branch lengths in Figure 1C also reveal that *Poseidon* and *Zeus* sequences rapidly diverged, in contrast to the extremely slow evolution of the parental *CAF40*. As an illustration, the CAF40 protein sequences from *D. melanogaster* and *D. willistoni*, which split around 36 MYA (Russo et al. 1995; Markow and O’Grady, 2007; Clark et al., 2007), diverged in only 7.7% of the sites in accordance with the extreme conservation in the protein sequences encoded by the *CAF40* homologues across all multi-cellular organisms (Fig. S1B), whereas *Poseidon* and *Zeus* diverged 48.6% and 22.5%, respectively, from *CAF40* in *D. melanogaster* (Table S1). An orthologous comparison of CAF40 and Poseidon protein sequences between *D. melanogaster* and *D. willistoni*, revealed an amino acid substitution rate of 0.12% per million years and 0.68% per million years, respectively. The comparison of *Zeus* protein sequence between *D. melanogaster* and *D. simulans,* which diverged by 3 MYA (Russo et al. 1995; Markow and O’Grady, 2007; Clark et al., 2007), obtains an unusually high substitution rate of 4.92% per million years.

### The duplicates diverged at highly conserved sites

In order to understand whether the duplicated proteins accumulated replacements at conserved residues in the ancestral protein, or merely at the highly variable termini of the protein, we estimated the Shannon entropy (H) for each residue in an alignment of *CAF40* homologs from 42 eukaryotes, and contrasted it with the replaced residues in the duplicates (Fig. S1). We found that amino acid replacements in Poseidon and Zeus occurred even at extremely conserved sites of *CAF40*. In both duplicates, replacements are distributed throughout the protein structure, including the charged groove formed by the conserved armadillo-repeat domain (Garces et al., 2007), which was shown to be important for *CAF40* interactions (Chen et al., 2014; Sgromo et al., 2017, 2018; Keskeny et al. 2019).

For the sake of comparison, out of the 49 completely conserved residues in CAF40 among eukaryotes (there are 49 conserved residues with which their H=0; Fig. S1C and S1D), Poseidon diverged in 25, and Zeus, in 11 of these completely conserved residues. Furthermore, several diverged residues that were experimentally shown to be functionally relevant for CAF40 interaction with its protein partners in previous studies were replaced in the duplicates (Table S2; Chen et al., 2014; Mathys et al., 2014; Sgromo et al., 2017). The extensive replacement of amino acids that are highly constrained in the parental protein suggests that Poseidon and Zeus functional properties may have diverged substantially from CAF40.

### The duplicates acquired a restricted expression pattern

Part of the phenotypic divergence between duplicated genes and their parents may result from their differential expression patterns. We used extensive transcriptome data from publicly available databases to investigate to which extent *Poseidon* and *Zeus* diverged from *CAF40* at the expression level. First, a comparison of expression of these genes in several tissues from *D. melanogaster* evidences that both *Poseidon* and *Zeus* have acquired a narrower expression pattern when compared to *CAF40* (Fig. 2A). The duplicates are only expressed at larval imaginal discs, adult male reproductive tissues, and pupae at low or intermediate levels, in sharp contrast with *CAF40*, which is expressed at all assayed tissues and development stages, from intermediate to high levels. We experimentally confirmed the distinct expression of the three paralogs in *D. melanogaster* larvae, head, and testis through RNA extraction followed by RT-PCR (Fig. S2).

**Figure 2.**
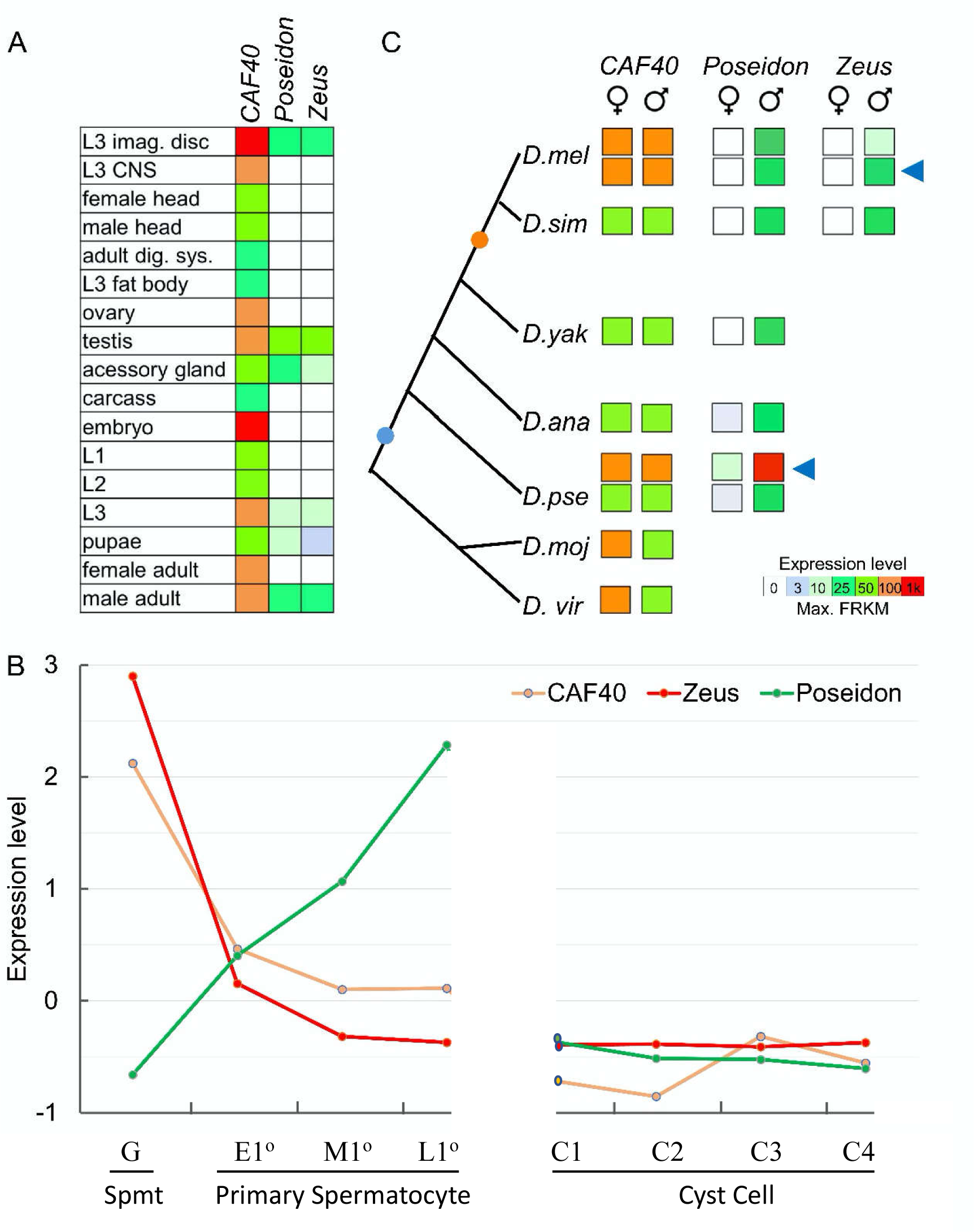
*CAF40*, *Poseidon* and *Zeus* have distinct expression patterns. A.) Summary of the expression intensity of each paralog in *D. melanogaster* tissues and development times. Expression summarized as average FPKM. *CAF40* broad expression contrasts with the duplicates restricted expression pattern. B.) Expression level, expressed as Z-scores of Transcriptions Per Kilobase Million (TPM) values of each gene to normalize by experimental variance (The Z-score of TPM normalized reads for each gene were directly retrieved from “Gene Level Data” Excel sheet in Supplementary Dataset 2 from Mahaderavaju et al., 2021), at eight different cell types in *D. melanogaster* testes (spermatogonia (Spmt, mitosis, G), primary spermatocyte (meiosis, early (E1°), middle (M1°) and late(L1°)), cyst cell differentiation (4 stages from C1 to C4)) (Mahadevaraju et al., 2021). These data provide evidence for the compensation of the autosomal *Poseidon* for the significantly reduced expression of the X-linked *CAF40* in the spermatocyte stages. Z-scores mean the number of standard deviations from the mean. Z = (x – μ)/σ, x is the value to be standardized, μ is the mean, σ is the standard deviation. C.) Summary of expression intensity in female/male adults for *Drosophila* species with available data for each paralog. Circles in the phylogeny represent the duplication event for *Poseidon* (blue) and *Zeus* (orange). Blue arrows indicate the expression intensity in reproductive organs (testes and ovaries) of *D. melanogaster* (*D. mel*) and *D. pseudoobscura* (*D. pse*). Male-biased expression for *Poseidon* and *Zeus* are conserved across the phylogeny. Data extracted from (Brown et al., 2014; VanKuren and Vibranovski, 2014; Vibranovski et al., 2009b).

We also compared the expression of each paralog at different testis cell types in *D. melanogaster*. We directly retrieved the Z-score of TPM normalized reads for each gene from a recently published dataset of genome expression during spermatogenesis using sing cell RNA-seq (scRNA-seq) from 8 spermatogenesis phases in mitosis, meiosis and cyst cell differentiation (see “Gene Level Data” Excel sheet in Supplementary Dataset 2 from Mahaderavaju *et al.,* 2021). We show that the three paralogs have distinct expression dynamics (Fig. 2B). The scRNA-seq detected expression patterns of the three genes are similar to those detected previously from dissected testis tissues (Vibranovski et al., 2009b), revealing the expression dynamics of these X-linked (*CAF40*) and autosomal genes (*Poseidon* and *Zeus*) in spermatogenesis with a much higher resolution. While *CAF40* expression level is high in spermatogonia, it reduces expression in meiotic and cyst cells, suggesting *CAF40* is negatively impacted by the inactivation of male X chromosome in spermatocytes. Zeus shows a strong peak of expression, higher than *CAF40*, in the early mitotic phase, and subsequent drop of expression in the meiotic and post-meiotic stages. Starting in a low level of expression in spermatogonia, *Poseidon* increases expression in the middle primary spermatocyte (M1°), eventually showing a peak of expression in the late primary spermatocyte (L1°), a common pattern for autosomal retrogenes expressed in the testis that compensates for the lowly expressed X-linked paralogues (Vibranovski et al., 2009b). Such a difference in expression is expected for retrogenes, which are inserted in genomic contexts diverse from the parental copy and may promptly acquire and/or evolve new cis-regulatory elements (Bai et al., 2008). The retroposed genes in autosomes may also help avoid the meiotic X chromosome inactivation (MXCI) when functioning in the meiosis stages of spermatogenesis (Betrán et al., 2002; Vibranovski et al., 2012; Mahaderavaju et al., 2020).

Finally, we analyzed transcriptome data from other six *Drosophila* species in order to understand whether the duplicates male-biased expression pattern is conserved across the phylogeny. In these six species, we found that *CAF40* is expressed at intermediate or high levels in adults, consistent with its role as a housekeeping gene. In contrast, these species exhibit significant male-biased expression of the duplicated genes (four additional species for *Poseidon*, and one for *Zeus*), similar to the pattern observed for *D. melanogaster* (Fig. 2C).

### *Poseidon* and *Zeus* impact viability and fertility

Restricted expression pattern of the duplicates, along with their conserved sex-specific expression across fly species, suggest that the duplicates may have been integrated into developmental and/or reproductive processes. First, we tested this hypothesis by using RNAi-mediated knockdown and CRISPR-based knockout to assay the functional effects of the three paralogs on *D. melanogaster* viability and fertility.

RNAi-knockdown using both a ubiquitous (*Tub84B*>GAL4) and an imaginal disc-specific driver (*T80*>GAL4) confirmed that *CAF40* expression is essential for survival. Less than 3% of the flies developed into adults when the gene was silenced with the ubiquitous driver (Fig. 3A, Fig. S3). An essential role of *CAF40* in cellular processes is also observed in distant eukaryotes, as evidenced by knockdown experiments in *C. elegans* (Kamath et al., 2003) and humans (Wang et al., 2015). Our knockdown assays detected a significant phenotypic impact on viability for *Poseidon* and *Zeus*. RNAi-knockdown of these genes reduced the relative fly viability by ∼20%, with a slightly stronger effect of the ubiquitous driver over the imaginal disc one (Fig. 3A). A more negative impact on viability was detected with ∼25% reduction when the genes were knocked out using CRISPR/Cas9 methods that created frameshift mutations by deleting 1 and 2 nucleotides in *Poseidon* and 4 and 8 nucleotides in *Zeus* (Fig. 3B). In comparison, an in-frame mutant by a deletion of 6 nucleotides in *Poseidon* did not have significant effect (Fig. 3B).

**Figure 3.**
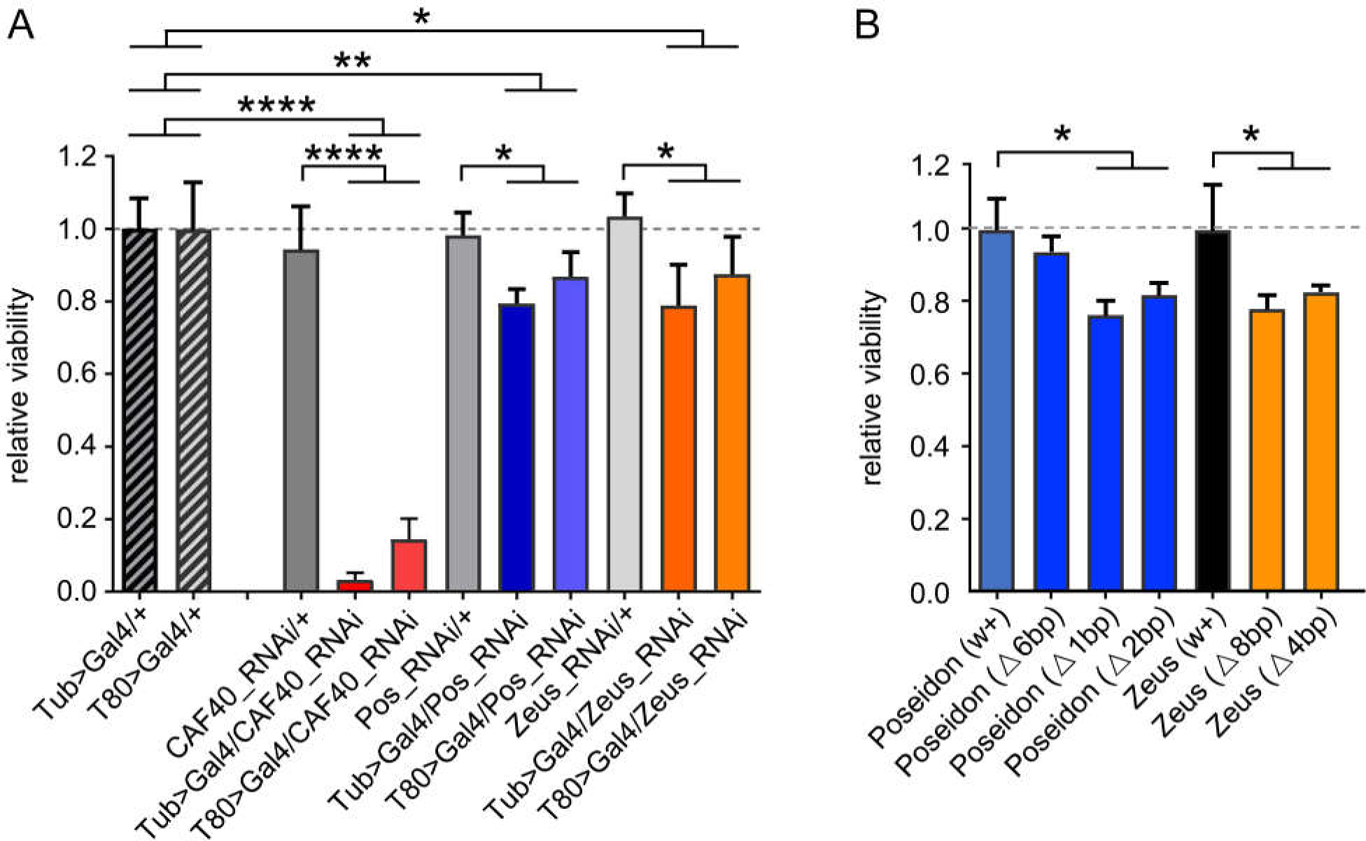
*CAF40*, *Poseidon* and *Zeus* impact on viability. A.) Viability of RNAi-expressing flies, relative to the control genotype from each individual cross. The two hatched bars in the left show the relative viability of the two control GAL4-expressing drivers, *Tub* (ubiquitous) and *T80* (imaginal-discs). *CAF40*-RNAi-expressing flies in red; *Poseidon* in blue and *Zeus* in orange. B.) Viability of homozygous flies for different CRISPR-Cas9 deletions for *Poseidon* (blue) and *Zeus* (orange) relative to the control from the same background strain (black). Notice the non-significant effect of the only non-frameshift deletion for *Poseidon* (first blue bar on the left); t-test: * p < 0.05, ** p < 0.01, *** p < 0.001, **** p < 0.0001. Error bars indicate SD.

Given the expression of the duplicates in testes, we further investigated the effect of these on male fertility. We used RNAi-knockdown with two different testis-specific drivers: *nanos*>GAL4 that silenced the gene expression at spermatogonia and male germline stem cells, and *Bam*>GAL4 that silenced gene expression at late spermatogonia and early spermatocytes stages (White-Cooper, 2012). These drivers allowed us to assay the fertility effects at different spermatogenesis phases. Independent silencing by the two drivers detected significant fertility effects in all three genes (Fig. 4). The *CAF40* knockdown showed the strongest effect (∼40% fertility reduction with *nanos*), in accordance with its function as a fundamental housekeeping gene. It’s worth noting that Chen et al. 2012 used a KK RNAi line of CAF40 (KK101462) with same driver and showed no significant fertility effect (See Materials and Method). The *nanos* driver also caused a 20% and 30% fertility decrease when knocking down *Poseidon* and *Zeus*, respectively. In contrast, the knockdown using the *Bam* driver caused the strongest fertility defect for *Poseidon* (30% fertility reduction) and a lower but significant effect for *CAF40* (20% fertility reduction), suggesting *CAF40* may play a more important role in an earlier stage (spermatogonia) of spermatogenesis than *Poseidon* with a more critical role in spermatocytes and/or later stages of spermatogenesis. *Zeus* appeared not to play any significant role in the later stages of spermatogenesis that was silenced by the *Bam*-driver (t-test, p = 0.59). Instead, *Zeus* was knocked down with a significant effect at the earlier stage by the *nanos* driver (Chen et al., 2012). These observations are in accordance with the expression pattern of the paralogs during spermatogenesis (Fig. 2).

**Figure 4.**
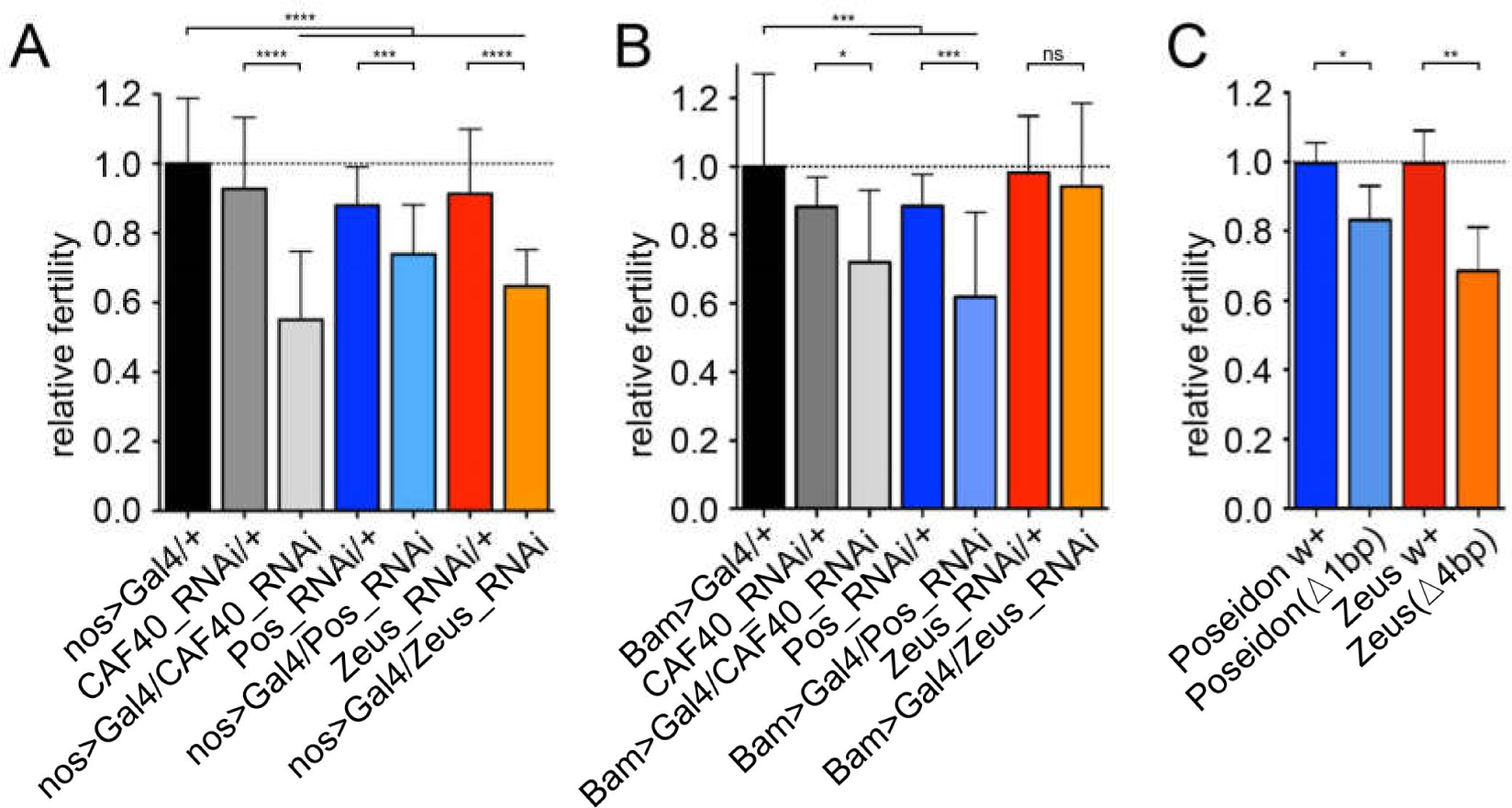
*CAF40*, *Poseidon* and *Zeus* impact on male fertility. Relative fertility of males expressing RNAi using testis-specific drivers: A.) *nanos*>GAL4 (early spermatogenesis); and B.) *Bam*>GAL4 (late spermatogenesis). Black bars represent the controls, *CAF40*-RNAi-expressing flies in grey, *Poseidon* in blue, *Zeus* in orange. C.) Fertility of males homozygous for frameshift CRISPR-Cas9 deletions for *Poseidon* (blue) and *Zeus* (orange) compared to the control of the same background strain (darker bars). t-test: * p < 0.05, ** p < 0.01, *** p < 0.001, **** p < 0.0001. Error bars indicate SD.

Finally, *Poseidon* and *Zeus* impact on male fertility was also confirmed by the knockout analyses using CRISPR/Cas9-generated frameshift deletions, significantly decreasing male fertility by 17% and 32%, respectively (Fig. 4C). In summary, a combination of knockdown and knockout analyses reveals that *Poseidon* and *Zeus* carry out important functions in support of viability and male fertility in the *Sophophora* subgenus (*Poseidon*) or the *D. melanogaster* subgroup (*Zeus*).

### Poseidon and Zeus interaction with the *CCR4-NOT* complex

CAF40 is a highly conserved subunit of the CCR4-NOT complex (Miller and Reese, 2012) and evolved slowly as a highly conserved gene in aforementioned analyses (Fig. 1). However, both Poseidon and Zeus have intense protein sequence divergence with CAF40. We then try to test whether Poseidon and Zeus, the two duplicates retroposed from CAF40, maintained its ancestral functions or evolved novel gene functions. First, we independently expressed a GFP-tagged version of each paralog in *Dm* S2 cells and assayed their interaction with HA-tagged NOT1 through co-immunoprecipitation followed by Western blotting analysis. NOT1 was selected because it is the central scaffold subunit of the CCR4-NOT complex and it was shown to directly interact with CAF40 via a CAF40/CNOT9-binding domain (CN9BD) (Collart and Panasenko, 2017; Mathys et al., 2014; Chen et al., 2014). Interestingly, we found that Poseidon maintained the ability to interact with NOT1, whereas Zeus either lost it, or binds NOT1 only weakly (Fig. 5A).

**Figure 5.**
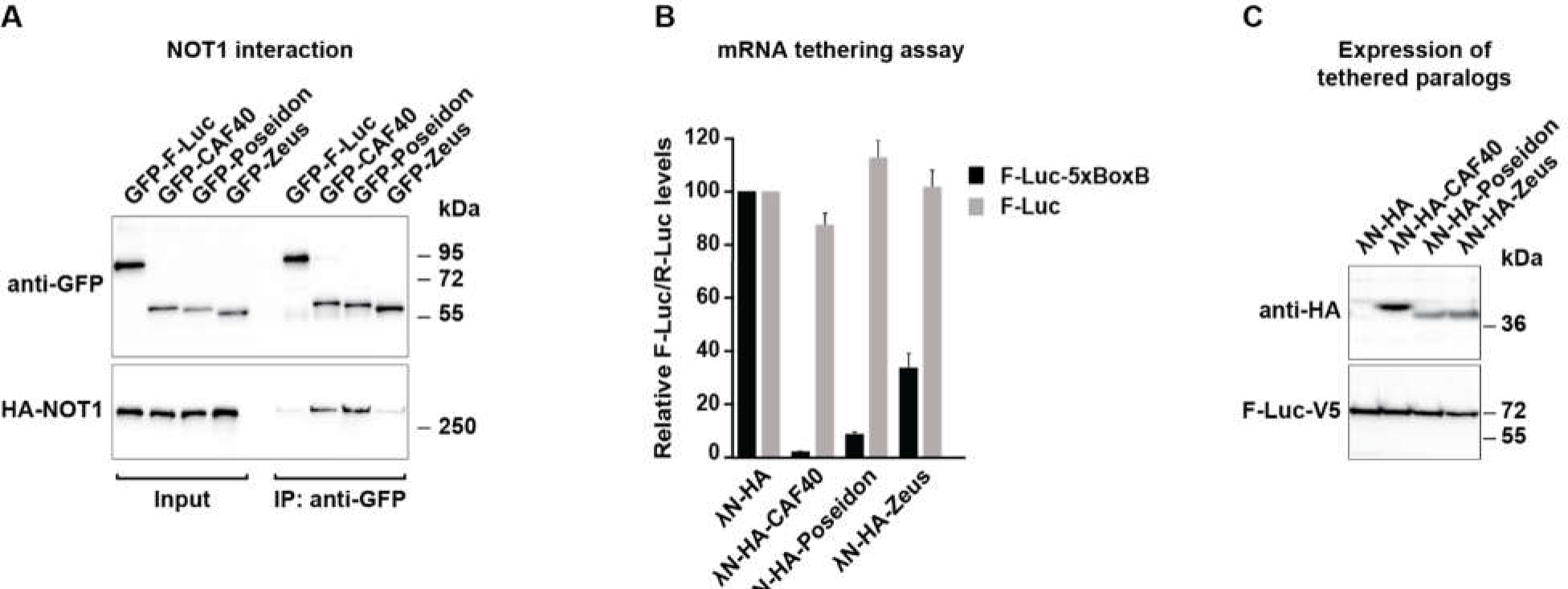
CAF40, Poseidon and Zeus protein interaction with the CCR4-NOT complex. A.) Co-immunoprecipitation assay showing the interaction of CAF40 paralogs with NOT1 in *Dm* S2 cells. Co-immunoprecipitation was conducted in the presence of RNase A to exclude RNA-mediated interactions. Cell lysates expressing GFP-tagged CAF40, Poseidon and Zeus, and HA-tagged NOT1. GFP-F-Luc served as a negative control. Input samples consist of 3% for the GFP-tagged proteins and 1% for the HA-tagged proteins, and immunoprecipitated samples of 10% for the GFP-tagged proteins and 30% for the HA-tagged proteins. Protein size markers are shown on the right in each panel. B.) Tethering assay using λN-HA-tagged CAF40, Poseidon and Zeus and the F-Luc-5BoxB reporter in *Dm* S2 cells (black bars). A plasmid expressing R-Luc served as a transfection control, and an F-Luc reporter lacking the 5BoxB binding sites for λN- was used as control (grey bars). F-Luc activity levels were normalized to those of the R-Luc control and set to 100% in cells expressing the λN- HA peptide alone. Error bars indicate SD of five replicates. CAF40 and Poseidon exhibit similar abilities of repressing the luciferase reporter (black bar) compared to the control (grey bar). Zeus exhibits lower, though still significant, repression ability. C.) Western blot analysis showing the equivalent expression of the λN-HA-tagged proteins used in the tethering assays shown in Figure 5B. Protein size markers (kDa) are shown on the right of the panel.

The conservation of the interaction with NOT1 observed for Poseidon suggests that it could be incorporated into the CCR4-NOT complex, in contrast to Zeus. We further infer that, if this is true, Poseidon should have conserved the repressive effect on targeted mRNAs observed for CAF40 (Bawankar et al., 2013; Sgromo et al., 2017). We tested this hypothesis by measuring each paralogs ability to repress a luciferase reporter mRNA in a λN/BoxB tethering assay. In the tethering assay the λN-tagged CAF40 paralogs are efficiently recruited to the reporter RNA, carrying five BoxB elements due to the strong binding of the λN peptide to the BoxB elements. In agreement with previous reports, tethering of λN-tagged CAF40 to a luciferase reporter mRNA carrying five BoxB elements in the 3′UTR (F-Luc-5xBoxB; black bar) strongly represses the protein synthesis of the reporter. In contrast, CAF40 did not affect the expression of the control F-Luc mRNA lacking BoxB elements (grey bar) (Fig. 5B). Intriguingly, Poseidon is also able to reduce the reporter expression to similar levels (around 10% of the control level, Fig. 5B), indicating that Poseidon can act together with post-transcriptional regulators such as the CCR4-NOT complex to modulate the luciferase mRNA. In contrast, Zeus exhibits a weaker repressive ability compared to the other paralogs, although it is clearly significant (around 40% of the control level). All paralogs were expressed at comparable levels (Fig. 5C).

Taken together, these results suggest that Poseidon conserved CAF40’s ability to interact with the CCR4-NOT complex through interactions with NOT1 most likely leading to the degradation of targeted transcripts. Zeus, however, lost or weakened its CCR4-NOT recruitment ability. Nevertheless, the fact that Zeus is still able to decrease reporter protein levels when tethered to an mRNA, suggests that it either evolved new protein interactions involved in mRNA regulation or that the weaker CCR4-NOT recruitment ability is sufficient to mediate repression in the tethering assay.

### *CAF40*, *Poseidon* and *Zeus* impact gene regulation

Given the central role of CAF40 in several cellular regulatory processes, we investigated the impact of the three paralogs on global gene expression. Moreover, since the two duplicates, Poseidon and Zeus, are highly diverged at their protein sequence and expression profile from the parental CAF40, they provide a suitable system to assay global expression regulation among the paralogs.

We conducted a genome-wide transcriptome analysis of adult testes to assay the impact of *CAF40*, *Poseidon* and *Zeus* on global gene expression using germline-specific knockdown with *nanos*-GAL4 drivers (Table S3, Fig. S4-S6). Our transcriptome data showed that RNAi-silencing was effective and specific for each paralog, reducing mRNA levels of each gene in at least 60% compared to the control, while not impacting the other paralogs (Table S4). The knockdown of each of the three genes detected over one thousand differentially expressed genes (DEGs) (Fig. 6A-6B, Table S5-S7). Putting together, the knockdowns of the three genes affected the expression of totally 2,622 genes (union set of different expressed genes from the three KD lines) (Fig. 6B), corresponding to more than a fifth of the genes mapped in our transcriptome (11,491) (Fig. S7-S8, Table S3). Such a widespread effect on gene expression suggests that *Poseidon* also plays an important role on gene regulation in spermatogenesis, as previously shown for *CAF40* and *Zeus* (Chen et al., 2012).

**Figure 6.**
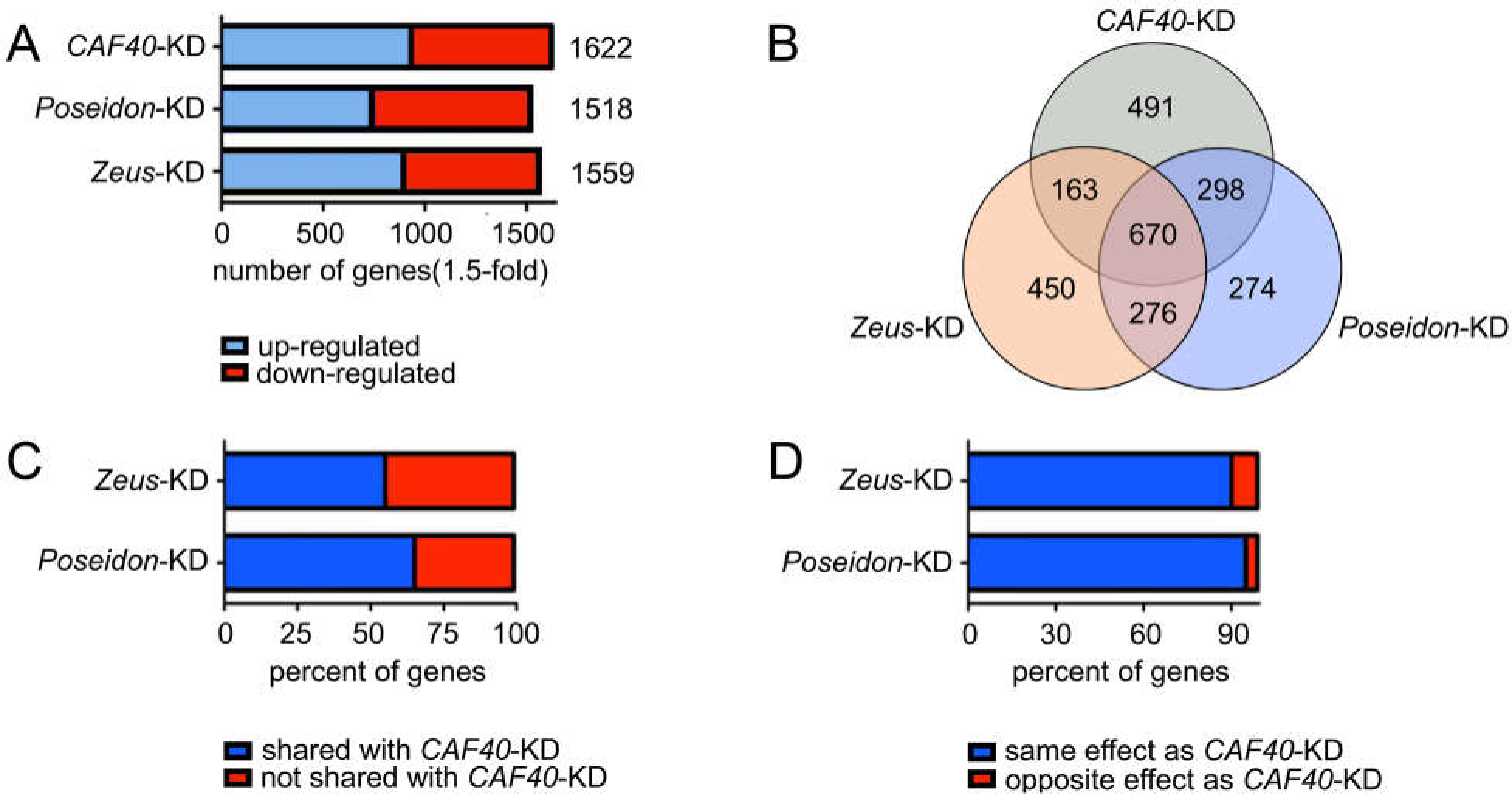
Impact of *CAF40*, *Poseidon* and *Zeus* knockdowns on global gene expression. A.) Number of genes with >1.5-fold change in expression compared to the control in the knockdown of each paralog measured through RNA-seq. Up- regulated genes shown in blue, down-regulated shown in red. B.) Venn diagram representing the number of genes differentially expressed upon the knockdown of each paralog (*CAF40* in grey, *Poseidon* in blue, *Zeus* in orange) and their intersections. Notice the large number of genes (670) commonly affected by the three knockdowns, as well as genes exclusively perturbed by one paralog (1,215). C.) Proportion of genes differentially expressed upon *Poseidon* and *Zeus* knockdown that are also affected by *CAF40* knockdown. Notice that *Poseidon* affects a higher proportion of genes shared with *CAF40* compared to *Zeus* (63.7% versus 53.3%; χ^2^ = 50.7, p < 10^-10^). D.) Proportion of genes differentially expressed upon the knockdown of *Poseidon* and *Zeus* that are affected in the same direction (i.e. up- or down- regulated) in the *CAF40*-KD. Notice that, despite the fact that the large majority of genes are affected in the same direction by the duplicates and the parental knockdown, *Zeus* knockdown affects a larger proportion of genes in the opposite direction of that of *CAF40* when compared to *Poseidon* (9.1% of perturbed genes in *Zeus*-KD versus 3.8% in *Poseidon*-KD were in the opposite direction of *CAF40*-KD, χ^2^ = 21.3, p < 10^-5^).

We also investigated the differential impact of these paralogs. We classified the genes with perturbed expression compared to the controls in each knockdown sample as male or female-biased, based on two independent *Drosophila* databases (Zhang et al. ,2010; Assis et al., 2012). KD of both *Zeus* and *Poseidon* shows bias towards both downregulating male-biased genes and upregulating female-biased genes (Table S8, χ2 test, p-value<0.05). These results indicate that both *Poseidon* and *Zeus* evolved function of activating male-biased genes expression and repressing female-biased genes expression (a consistent result with Chen et al., 2012). However, we couldn’t conclude here that the upregulated female-biased genes of Zeus-KD were significantly enriched on the X chromosome (Table S8, χ2 test, p-value=0.4431). This may be caused by the very low number of the intersection genes between X-linked female-biased genes and those down/up- regulated genes of the three KDs (Table S8). However, the downregulated male-biased genes of both Poseidon-KD and Zeus-KD were overrepresented on autosomes (Table S8, χ2 test, both p-value<0.01), consistent with previously reported chromosomal distribution patterns of sex-biased genes(Chen et al., 2012).

A large set of 670 genes (25.5% of total 2,622 impacted genes) was perturbed when any of the three paralogs was individually silenced (Fig. 6B). Interestingly, the cellular processes with the most significant enrichment in these genes were proteolysis (GO:0006508, adj. p < 10-7, both of the results by the tools Gorilla and g:Profiler confirm this cellular processes) and reproduction (GO:0032504, adj. p < 10^-7^) (Table S9). *Poseidon* shares the most genes with *CAF40*, in comparison to other two two-gene comparisons (298 versus 163, 276, respectively) whereas *Zeus* shared least number (163) of genes with *CAF40* (Fig. 6B). Furthermore, in the shared genes with *CAF40*, *Poseidon* has a significantly higher proportion of genes in the same direction towards up- or down-regulation whereas Zeus has opposite changes, e.g. a gene with up-regulation for *CAF40* and down-regulation for *Zeus* (Fig. 6C-6D, Table S10) (Chi-squared test, p=0.000699, <0.01). These observations are in accordance to the finding that *Poseidon* and *CAF40* behave more similarly in regard to protein-protein interactions and repressive activity in comparison to *Zeus* (Fig. 5).

Nevertheless, a substantial set of genes (1,215 genes, 46.3%) was perturbed by only one of the knockdowns, but not shared with the other two, which reveals the distinct impact that each paralog has in the global regulatory network (Fig. 6B). This suggests that three paralogues have evolved peculiar interactions with large numbers of non-overlapping genes. Interestingly, Poseidon has the least number of such peculiar gene interactions (274 versus 491 (*CAF40*), 450 (*Zeus*), respectively). Additionally, we calculated the DEGs intersection numbers of those subgroups between our study and Chen et al., 2012 (Microarray) and found those DEGs intersection numbers between two studies are all relatively small. This may be caused by different experiments conditions (See Materials and Method).

## DISCUSSION

We showed that a functionally important and conserved member of the CCR4- NOT complex, CAF40, gave rise to two gene duplicates through retroposition, *Poseidon* and *Zeus,* in recent evolution of *Drosophila* species. We demonstrated that *Poseidon* and *Zeus* are functionally important genes that have quickly diverged from *CAF40* in protein sequence and expression shortly after duplication, whereas the parental *CAF40* remained highly conserved (Fig. 1C). Remarkably, even residues that have been conserved in *CAF40* for a long evolutionary time (e.g. amino acids identical in all eukaryotic homologs) were extensively substituted in the duplicates, which may impact conserved functions of the protein (Fig. S1).

However, our molecular functional analyses show that both *Poseidon* and *Zeus* are important spermatogenesis genes. *Poseidon* retained mRNA suppression functions of *CAF40* while *Zeus* evolved divergent functions as a suppressor of female genes in males (Chen et al., 2012).

Our co-immunoprecipitation assay showed that Poseidon protein conserved CAF40 ability to interact with NOT1. Moreover, such interaction is consistent with CAF40 and Poseidon, both showing a strong repressive effect on a tethered reporter transcript (Fig. 5). Zeus retains a lower repressive activity, which probably reflects its divergent ability on acting in protein stability. These data suggest that, whereas Poseidon likely has inherited the CAF40 role in the CCR4-NOT complex (although with a distinct impact on gene regulation, as discussed below), Zeus likely functions independently of this complex as a suppressor of femininized genes (Wu and Xu, 2003), as the genomic DNA binding has shown, revealed by ChIP-chip analysis (Chen et al., 2012).

Given the high expression and the conserved functions of *CAF40* in post-transcriptional gene regulation, an important question is raised: why did the *Sophophora* subgenus evolve an additional copy, *Poseidon*, to encode a similar function in mRNA regulation? The duplications of the X-linked CAF40 in form of the autosomal Poseidon and Zeus are likely a consequence of natural selection acting to comply with MXCI in evolution of sex chromosomes in *Drosophila* (Mahadevaraju et al., 2021; Vibranovski et al., 2009b; Betrán et al., 2002). An autosomal location can help these new genes to play their functional roles by escaping expression suppression by MXCI as its X-linked CAF40 experiences during inactivation. The fitness effects that these new gene duplicates brought under natural selection were detected to be critically important in viability during development and male specific fertility. Given the divergence of *Zeus* from *Poseidon*, these data suggest likely different roles of the two paralogs. *Poseidon* may compensate for *CAF40* whose expression is suppressed by MXCI, similar to the function of the autosomal retroposed *RPL10L* in humans (Jiang et al., 2017. Long and Emerson, 2017). However, *Zeus* is not completely redundant to *Poseidon*. The divergence in DNA sequence and expression make them functionally distinct in spermatogenesis. Additionally, a more complicated model was the SAXI model of the sexual antagonistic selection on the sex chromosome (Wu and Xu, 2003). The model argued that the selection leading to demasculinization of X chromosomes before the establishment of the silencing of X chromosome (or regions) in evolution can also pressure a X->A gene traffic through duplication including retroposition (Emerson et al, 2004). Related to this, female germlines, which are not subject to the X inactivation in Drosophila, may also express in a lower level the X-linked male genes that are antagonistically selected against. These genes in females can serve as substrate for retroposition as well.

Our genome-wide transcriptome analyses demonstrated that the independent perturbation of the three paralogs impacts the regulation of thousands of genes in the testes (Fig. 6). This is in agreement with the important role of CAF40 in transcriptional and post-transcriptional regulation (Collart, 2016), as well as with the significant role of *Zeus* as a suppressor of female genes in males (Chen et al., 2012). The analysis of the genes that are commonly perturbed by the paralogs’ knockdown compared to the control (Fig. 6B), revealed a strong enrichment for genes related to catabolic function (Table S9), such as serine-type endopeptidase activity (GO:0004252, p<10-28), serine hydrolase activity (GO:0017171, p<10-27), and catalytic activity (GO:0003824, p<10-6). Those enrichment suggests that the knockdown of the paralogs affects the regulatory balance between transcription, translation and degradation of numerous downstream genes, given the importance of the parental gene in coordinating and integrating different regulatory pathways (Miller and Reese, 2012).

Taken together, the analyses presented here suggest that CCR4-NOT, a multifunctional complex that controls gene expression at multiple levels within the cell, evolved a new member *Poseidon*, through retroposition from *CAF40*, in *Drosophila*. *Poseidon* and *Zeus*, despite their relatively recent origination in *Drosophila*, were integrated into fundamental cellular and molecular processes with profound impacts in the regulatory network and phenotype. They were selected for compensation for the inactivated X-linked *CAF40* during male meiosis or unrelated new functions, respectively. They reveal that a fundamentally important and conserved gene function also evolved with quick gene evolution, driven by evolution of sex chromosomes with its ancestral generated MXCI.

## MATERIALS AND METHODS

### Molecular evolutionary analyses

*Poseidon* (*CG2053*) had been previously computationally identified as a putative young gene (Zhang et al., 2010). Gene and protein sequences were retrieved from Flybase and NCBI, aligned with MUSCLE (Edgar, 2004) and manually curated. Reciprocal PSI-BLAST (NCBI) searches were employed to survey for *CAF40*, *Poseidon* and *Zeus* orthologs in eukaryotes. The proper substitution model for the alignment (GTR+G) was selected through a likelihood ratio test using jModeltTest (Posada, 2008). The phylogenetic relationship among the paralogs was firstly inferred through Bayesian analysis in MrBayes (Ronquist et al., 2012) by putting the three genes together. MCMC analysis was run with 4 chains for 2 million generations, with trees begin sampled every 500 generations, and the first 25% of samples were discarded as burn-in (Fig. S9). However, we found that there is no way to avoid long branch attraction (LBA) when invoking both rapidly evolving genes, *Zeus* and *Poseidon*, or branch length attraction (BLA) when invoking extremely slowly evolving *CAF40* and the two rapidly evolving genes to generate congruent trees between genes tree and species three. We have also tried the three classical ways (ML, NJ and MP) to construct three phylogenetic trees by putting the all the orthologs of the three paralogs together, and find they have same problem as Fig. S9. However, all those three phylogenetic trees indicated that the Zeus orthologs’ cluster are always closer to CAF40 orthologs’ cluster than Poseidon. This indicates that Zeus originated from CAF40 (Bai et al., 2007; Quezada-Díaz et al., 2010). Finally, to avoid LBA and BLA effect, we respectively constructed the congruent phylogenetic trees using the classical maximum parsimonious method by separating the three paralogs (Fig. 1C).

For the divergence of expression analysis, we retrieved the summary of expression data (FPKM [fragments per kilobase per million mapped reads]) from modENCODE and public RNA sequencing data of diverse fly species (Brown et al., 2014; Chen et al., 2014; VanKuren and Vibranovski, 2014). Expression values of each paralog at different spermatogenesis stages in *D. melanogaster* was compared using data from the SpermPress database (Vibranovski et al., 2009b).

### Shannon Entropy analyses

Shannon’s entropy was calculated for an alignment of CAF40 orthologous protei n sequences from 56 eukaryotes (Fig. S1), and the entropy value (H) for each residu e was plotted onto CAF40 protein structure from D. melanogaster (Sgromo et al., 20 17) using PyMOL. We used Shannon entropy (H) (Shannon, 1948), with calculation of H score which represents standard entropy for a 22-letter alphabet. The calculatio n by bio3d follows: http://thegrantlab.org/bio3d/, or https://bitbucket.org/Grantlab/bio3 d/downloads/, https://github.com/Grantlab/bio3d.git, bio3d_2.2-2.tar.gz, last-accesse d date: May 2019. (Grant, 2006; Mirny and Shakhnovich, 1999).

### Knockdown and knockout phenotypes

In order to assay the knockdown effect of each paralog on egg to adult viability, homozygous UAS-TRiP RNAi lines (Perkins et al., 2015) were crossed to a balanced constitutive driver line (*Tub84B*>GAL4/TM3) and an imaginal disc-specific driver line (*T80*>GAL4/CyO) (Table S11 shows the list of lines used). At least 10 independent replicates of 3 couples were allowed to cross and lay eggs for 7 days at 23°C. All F1 adults in the progeny were scored, and the proportion of wild/balancer phenotypic markers for all replicates was compared to control crosses (TRiP background line BDSC 36303 crossed to the driver lines). Male fertility effects were assayed for the three paralogs by driving GAL4 expression using male-germline-specific *nanos*- GAL4 and *Bam*-GAL4 drivers, which are expressed in early and late spermatogenesis, respectively (Table S11).

Of particular note is that Chen et al (2012) used a RNAi line (KK101462) of CAF40 from KK (phiC31) RNAi line library in VDRC (https://stockcenter.vdrc.at/control/library_rnai). Based on our lab empirical data, the KK library may have a lower knockdown efficiency than GD library (Xia et al., 2021), we tried to seek other mutant to reanalysis the effect of this gene. Finally, we used the UAS-TRiP RNAi lines in this study. TRiP uses a short but specific dsRNA (∼21bp), while KK uses a longer dsRNA (81-799bp, average:357bp) and they used different vector with varying KD efficiency (KK: pkC26; TRiP: pVALIUM20). Ni et al., (2011) found that the VALIUM20 (TRiP line construction vector) gives a stronger knockdown than the long-hairpin–based vector VALIUM10 in the soma and works well in the germline. This may cause the different phenotype effect of *CAF40* between our study and Chen et al., 2012. The second potential reason may be the different target site of the two lines: KK101462 target the last exon and the 3’UTR region of *CAF40* while TRIP.HMS05850 target to the third exon of *CAF40*. Therefore, we believe that dsRNA vector and target position may cause the different phenotypes of one same gene.

At least 15 replicates with 3-5 days old knockdown males were individually crossed to two virgin females from background line BDSC 36303 for one day. Females were allowed to lay eggs for 7 days, and all the F1 adults were counted. Knockdown efficiency for each paralog was well confirmed (the expression was reduced to 15∼35% of the control) through RT-PCR (Figure S3). RNA samples were extracted in triplicate using RNeasy kit (Qiagen, Cat. No. 74104), digested with DNase I (Invitrogen, Cat. No. 18047019) to remove genomic DNA contamination, and reverse transcribed with SuperScript III Reverse Transcriptase (Invitrogen, Cat.

No. 18080093) using oligo(dT) primers. RT-PCR was performed using iTaq Universal SYBR Green Supermix (Bio-rad, Cat. No. 1725121), with three technical replicates for each biological replicate. Quantitative PCR values were normalized using the ΔΔCT method to the *Rp49* control product.

CRISPR-Cas9 frameshift deletions were induced for *Poseidon* and *Zeus*. Guide RNAs were designed using CRISPR Optimal Target Finder (Table S12, http://targetfinder.flycrispr.neuro.brown.edu/) to target early portions of the exon, and injected (300 ng/uL) along with Cas9 protein (PNA Bio Lab: CP01, 500 ng/uL) into embryos from the BDSC 25710 line (Bassett and Liu, 2014). F1 mutant individuals were screened and crossed to balancer lines (w^+^;Sb/TM3; and w^+^;Sco/CyO, respectively). Small frameshift deletions were confirmed through Sanger sequencing and created early stop codons in the transcribed genes (Fig. S10). Viability and male fertility assays were performed with knockout flies as described above, using the injected line BDSC 25710 as control.

### Co-immunoprecipitation assay

DNA constructs with the coding region of *D. melanogaster* genes *CAF40*, *NOT1*, used were described before (Sgromo et al., 2017, 2018). Plasmids encoding Poseidon and Zeus were generated by inserting the corresponding cDNA (Thermo Scientific) into the pAc5.1-λN-HA and pAc5.1-GFP vectors (Rehwinkel et al., 2005; Tritschler et al., 2007) using HindIII and XhoI restriction sites. All constructs were confirmed by Sanger sequencing. For the co-immunoprecipitation assay in *D. melanogaster* S2 cells (ATCC), 2.5×10^6^ cells were seeded per well in 6-well plates and transfected using Effectene transfection reagent (Qiagen, Cat. No. 301425). The transfection mixtures in Figure 5A contained 1 μg, 1.8 μg and 1.8 μg of plasmids expressing GFP-tagged CAF40, Poseidon and Zeus, respectively. 1 μg of HA-tagged NOT1 was used.

Cells were harvested 3 days after transfection, and co-immunoprecipitation assays were performed using RIPA buffer [20 mM HEPES (pH 7.6), 150 mM NaCl, 2.5 mM MgCl2, 1% NP-40, 1% sodium deoxycholate supplemented with protease inhibitors as previously described (Tritschler et al., 2008; Sgromo et al., 2018)]. All co-immunoprecipitation assays in S2 cell lysates were performed in the presence of RNase A as previously described (Sgromo et al., 2017). All Western blots were developed using an ECL western blotting detection system (GE Healthcare, RPN2232). The antibodies used in this study are listed in Table S13.

### Luciferase assay

For the λN-tethering assays in *D. melanogaster* S2 cells, 2.5×10^6^ cells per well were seeded in six-well plates and transfected using Effectene transfection reagent (Qiagen, Cat. No. 301425). The transfection mixtures contained the following plasmids: 0.1 μg of Firefly luciferase reporters (F-Luc-5BoxB or F-Luc-V5), 0.4 μg of the Renilla Luciferase (R-luc) transfection control and various amounts of plasmids expressing the λN-HA-tagged paralogs (0.01 μg for CAF40, 0.1 μg for Poseidon, 0.02 μg for Zeus). The plasmids for tethering assays in S2 cells (F-Luc-5BoxB, F- Luc-V5 and R-Luc) were previously described (Behm-Ansmant et al., 2006; Zekri et al., 2013). Cells were harvested 3 days after transfection and Firefly and Renilla luciferase activities were measured by using a Dual-Luciferase Reporter Assay System (Promega, Cat. No. E1910). The mean values +/- SD from five independent experiments are shown.

### RNA-seq analysis

Total RNA was extracted with Arcturus^TM^ PicoPure^TM^ RNA Isolation kit (Applied Biosystems, LOT 00665884) from testes of 3-5 days old knockdown males and controls, with three biological replicates. A total amount of 1 μg RNA per sample was used to construct the cDNA library, using NEBNext Ultra RNA Library Prep Kit for Illumina (NEB, #E7770) following manufacturer’s recommendations. Briefly, poly(A) mRNA was purified from total RNA using oligo(dT)-attached magnetic beads, reverse-transcribed to double-stranded cDNA with random primers, end-repaired and ligated with NEB adaptors for Illumina, before sequencing (HiSeq 4000, University of Chicago Genomics Core Facility).

Raw reads were processed and mapped to *D. melanogaster* reference genome (dm6) using STAR with default parameters (Dobin et al., 2013), and evaluation of transcriptional expression was carried out using featuresCounts (Liao et al., 2014).

For the differential expression analysis, methods DESeq2 (Love et al., 2014, v1.21.22), edgeR (Robinson et al., 2010, v3.23.5), and limma (Ritchie et al., 2015, v3.37.7) were independently employed. Genes were considered as differentially expressed if they were consensually called by the three methods (Fig. S11), with an expression fold change of at least 1.5 compared to the control at false discovery rate less than 0.1. For differentially expressed genes, enriched biological processes and molecular functions were identified using both GOrilla (Eden et al., 2009) and g:Profiler (Raudvere et al., 2019), with p-values < 10^-4^, and a false discovery rate of 0.1. Both of the two tools showed same result. The interaction network of proteins with enrichment for catabolic processes was visualized using STRING (von Mering et al., 2003), selecting only experimentally validated interactions with high confidence. The analyses of differentially expressed genes with male/female-biased expression followed two independent *Drosophila* databases (Zhang et al. ,2010; Assis et al., 2012).

We calculated the DEGs intersection numbers of those subgroups between our study and Chen et al., 2012 (Microarray). For example, only 241/1784=13.5% genes out of *CAF40*_DEGs_Chen was overlapped with *CAF40*_DEGs_Xia. The intersection between those 833 genes (intersection between *CAF40*_DEGs_Xia and *Zeus*_DEGs_Xia) and 664 genes (intersection between *CAF40*_DEGs_Chen and *Zeus*_DEGs_Chen) is only 50 genes (Table S10). Also, only 36 genes were overlapped between *CAF40*_downregulated DEGs in Chen et al. 2012 and *CAF40*_downregulated DEGs in our study. Those small intersection numbers between our study and Chen et al., 2012 were mainly caused by different approaches, RNAi lines and developmental stages: firstly, Chen et al., 2012 used GD49820 and KK101462 as *Zeus* and *CAF40* RNAi lines, respectively. However, our study used TRiP RNAi lines for both *Zeus* and *CAF40*. Secondly, Chen et al., 2012 applied microarray expression profiling to obtain the DEGs of *Zeus* and *CAF40* KD while our study used RNA-seq. Thirdly, Chen et al., 2012 used testis from 1-7 days old males while our study used testes of 3-5 days old knockdown males. Therefore, even those small intersection numbers between our study and Chen et al., 2012, among the DEGs in *Zeus* and *CAF40* knockdown genotypes, our study also showed less than 50% DEGs show changes in the same direction (46.6% and 48.5%, Tables S10). Additionally, *Poseidon* knockdown genotypes showed a slightly higher proportion of DEGs (57.3% and 61.2%, Tables S10) which showed changes in the same direction with *CAF40*.

## SUPPLEMENTARY FIGURE LEGENDS

**Figure S1. The divergence and conservation of protein sequences of CAF40, Poseidon and Zeus.** Alignment of CAF40, Poseidon and Zeus showing their high level of divergence in protein sequences (A.), in contrast to the extreme conservation of CAF40 across multicellular organisms, as shown in the alignment among an exemplified part of CAF40 orthologs from 42 eukaryotes (B.). Parameters for the sequence comparisons in A.: 1. Zeus versus Poseidon: Length: 302; Similarity: 156/302 (51.7%); Gaps: 14/302 (4.6%); Score: 340.0. 2. Zeus versus CAF40: Length: 306; Similarity: 232/306 (75.8%); Gaps: 16/306 (5.2%); Score: 982.5. 3. Poseidon versus CAF40: Length: 326; Similarity: 175/326 (53.7%); Gaps: 48/326 (14.7%); Score: 457.5. Shannon’s entropy (H) for each residue in the CAF40 alignment from eukaryotic orthologs (C.) and their frequency distribution as shown in the histogram plot of entropy values calculated for CAF40 (D.). The two bars on the left of D. represent very low H values, i.e. highly conserved residues in the alignment.

**Figure S2. Agarose gels confirming the expression of each paralog at different tissues from *D. melanogaster*.** RNA extraction was followed by DNAase treatment and reverse transcribed for each sample. Primers for *actin5C* were used as positive control. Small differences between the size of reverse-transcribed and gDNA bands are due to the presence of introns in *CAF40* and *actin5C*. A.) *CAF40* and *Poseidon*; B.) *Zeus* and *actin5C* as control. C- is a negative control.

**Figure S3. Constitutive RNAi-knockdown efficiency measured through quantitative PCR.** Expression levels of each target paralog relative to the normalized control (+/Tub>GAL4 driver), in larvae with the indicated genotypes. Quantitative PCR results were normalized using the ΔΔCT method to Rp49 product. Bars show means and SD for three replicates.

**Figure S4. MA plot of DEGs between Poseidon-KD and CAF40-KD line by deseq2.** The MA plot visualizes the differences between measurements taken in two samples, by transforming the data onto M (log ratio) and A (mean average) scales, then plotting these values. The CG name and gene symbol of those genes showing both more significant fold change and higher mean expression in Poseidon-KD compared to in CAF40-KD are indicated in MA plot figure.

**Figure S5. MA plot of DEGs between Poseidon-KD and Zeus-KD line by deseq2.** The MA plot visualizes the differences between measurements taken in two samples, by transforming the data onto M (log ratio) and A (mean average) scales, then plotting these values. The CG name and gene symbol of those genes showing both more significant fold change and higher mean expression in Poseidon-KD compared to in Zeus-KD are indicated in MA plot figure.

**Figure S6. MA plot of DEGs between Zeus-KD and CAF40-KD line by deseq2.** The MA plot visualizes the differences between measurements taken in two samples, by transforming the data onto M (log ratio) and A (mean average) scales, then plotting these values. The CG name and gene symbol of those genes showing both more significant fold change and higher mean expression in Zeus-KD compared to in CAF40-KD are indicated in MA plot figure.

**Figure S7. Clustering of samples upon knockdown in the RNA-seq assay.** A.) PCA plot of all the replicates in the control (green), *CAF40*-KD (grey), *Poseidon*-KD (blue) and *Zeus*-KD (orange). The x-axis represents PC1, and y-axis, PC2. B.) Heatmap showing normalized expression values of each sample for selected genes upon *CAF40*, *Poseidon* and *Zeus* independent knockdown, compared to the control. Each row represents a different gene, selected among all the differentially expressed ones, and each column represents a different replicate. Top panel: genes commonly affected by the knockdown of the three paralogs relative to the control; middle panel: genes only affected by *CAF40* knockdown; bottom panel: genes only affected by *Zeus* knockdown.

**Figure S8. Impact of *CAF40*, *Poseidon* and *Zeus* knockdowns on global gene expression.** Number of genes with >2-fold change (A.) and >5-fold change (B.) in expression compared to the control in the knockdown of each paralog measured through RNA-seq. Up-regulated genes shown in blue, down-regulated shown in red. C. Proportion of genes differentially expressed upon *Poseidon* and *Zeus* knockdown that are also impacted by *CAF40*. Notice that *Poseidon* affects a higher proportion of genes shared with *CAF40* at both 2-fold and 5-fold, compared to *Zeus*.

**Figure S9. Phylogenetic relationship among the three paralogs reconstructed through Bayesian method**. *CAF40* homologs in black, *Poseidon* in blue, and *Zeus* in orange; numbers represent the Bayesian posterior probability of each node.

**Figure S10. Knockout mutants for *Poseidon* (A) and *Zeus* (B) generated through CRISPR-Cas9.** Top panel: wild-type and mutant gene sequences for both duplicates confirmed through Sanger-sequencing. Bottom panel: sequencing chromatogram depicting a wild-type (top) and a heterozygous mutant (bottom). Red triangle represents the expected cut site. Notice the multiple peaks downstream of the cut sites, reflecting heterozygous alleles.

**Figure S11. DEGs analysis of CAF40-KD vs Control, Poseidon-KD vs Control and Zeus-KD vs Control by using edger, deseq2, and limma.** There are 1622 overlap DEGs between CAF40-KD and Control (left), 1518 overlap DEGs between Poseidon-KD and Control (middle), 1559 overlap DEGs between Zeus-KD and Control (right), by using the three commonly used tools (DESeq2, edgeR, and limma). Also see Figure 6A.

## SUPPLEMENTARY TABLE LEGENDS

**Table S1. The degree of sequence identity and similarity among the three paralogs in D. melanogaster.** This identity and similarity analysis were based on default parameters on https://www.ebi.ac.uk/Tools/psa/emboss_needle/.

**Table S2. Poseidon and Zeus accumulated mutations at functionally important residues in CAF40.** Below, replaced residues in Poseidon and/or Zeus proteins that were experimentally assayed in previous studies, and shown to be important for CAF40 function (Chen et al., 2012; Mathys et al., 2014; Sgromo et al., 2017).

Table S3. Global genes expression level and DEGs for all genotypes.

**Table S4. RNAi-knockdown efficiency for reducing the expression of each paralog in our RNA-seq assays**. Note that each knockdown significantly reduced the expression of its targeted gene, while not affecting the expression of the other two paralogs.

Table S5. Overlap gene list of DEGS from KD-CAF40.vs.KD-Ctrl by using three tools DESeq2, edgeR, and limma.

**Table S6. Overlap gene list of DEGS from KD-Poseidon.vs.KD-Ctrl by using three tools DESeq2, edgeR, and limma.**

Table S7**. Overlap gene list of DEGS from KD-Zeus.vs.KD-Ctrl by using three tools DESeq2, edgeR, and limma.**

**Table S8.** Proportion of differentially expressed genes (DEG) upon knockdown of CAF40, Poseidon and Zeus that exhibit significant either female- or male- biased expression according to two different databases (Assis et al., 2012; Zhang et al. ,2010). Notice that the knockdown of the three paralogs affects a significantly higher proportion of male-biased genes when compared to the total number of genes in our dataset.

Table S9**. List of Gene Ontology terms with the top 10 most significant enrichment among the DEGs upon *CAF40*, *Poseidon* and *Zeus* knockdown according to our RNA-seq analysis**. Only genes with p-values lower than 10^-4^ and false discovery rate q-value lower than 0.05 are shown (N=total number of genes; B=total number of genes associated with a GO term; n=number of genes with differential expression; b=number of genes in the intersection).

**Table S10.** Intersection of upregulated/downregulated genes between Poseidon Zeus and CAF40 in this study and Chen et al. 2012

Table S11**. List of fly strains used in this study.**

Table S12**. List of primers used in this study.**

**Table S13. List of antibodies used in the co-immunoprecipitation assays.**

## ACKNOWLEDGMENT

We dedicated this paper to Elisa Izaurralde, to show our deep respect for her devotedness to science and warm collegiality, besides her monumental contribution to the investigation of post-transcriptional and translational regulation. We remember that, in her last months, she enthusiastically arranged a fruitful collaboration with the Chicago team. She gave insightful discussion and designed with A.S. the experiments related to understanding the molecular function and evolution of Poseidon. We also appreciate Nicholas VanKuren’s constructive discussion with several details in our study. I. V. identified existence of *Poseidon* in the *Sophophora subgenus* of *Drosophila*. S. X. created the CRISPR mutants in this study. I. V., S. X., and S. H. conducted analyses of fitness effects, expression and evolution of *Poseidon* and *Zeus*. S. X. conducted phylogeny analyses, sequence and RNA-seq analyses. A. S., A. B. with E. I., investigated the molecular functions of Poseidon and Zeus. Wen-Qing Chan at the University of Chicago Center for Research Informatics provided valuable help in the RNA-seq analyses. M. L. and I. V. conceived the study of the new gene systems, M. L., I. V., S. X. and A. S. composed the manuscript. I.V. was supported by the Science without Borders Scholarship (BEX18816/12-6). M. Long was supported by NSF1026200 and NIH R01GM116113.

## Data Availability Statements

The data underlying this article are available in the article and in its online supplementary material.

## REFERENCES

1. Abrusan, G. (2013). Integration of New Genes into Cellular Networks, and Their Structural Maturation. Genetics 195, 1407–1417.

2. Assis, R., Zhou, Q., and Bachtrog, D. (2012). Sex-Biased Transcriptome Evolution in Drosophila. Genome Biol Evol 4, 1189–1200.

3. Bai, Y.S., Casola, C., Feschotte, C., and Betran, E. (2007). Comparative genomics reveals a constant rate of origination and convergent acquisition of functional retrogenes in Drosophila. Genome Biol 8, 1–9.

4. Bai, Y.S., Casola, C., and Betran, E. (2008). Evolutionary origin of regulatory regions of retrogenes in Drosophila. BMC Genomics 9, 1–9.

5. Baker, C.C., and Fuller, M.T. (2007). Translational control of meiotic cell cycle progression and spermatid differentiation in male germ cells by a novel eIF4G homolog. Development 134, 2863–2869.

6. Bawankar, P., Loh, B., Wohlbold, L.,Schmidt, S. & Izaurralde, E. (2013). NOT10 and C2orf29/NOT11 form a conserved module of the CCR4-NOTcomplex that docks onto the NOT1 N-terminal domain, RNA Biology 10, 228–244.

7. Bassett, A., and Liu, J.L. (2014). CRISPR/Cas9 mediated genome engineering in Drosophila. Methods 69, 128–136.

8. Behm-Ansmant, I., Rehwinkel, J., Doerks, T., Stark, A., Bork, P., and Izaurralde, E. (2006). MRNA degradation by miRNAs and GW182 requires both CCR4 : NOT deadenylase and DCP1 : DCP2 decapping complexes. Genes Dev 20, 1885–1898.

9. Belote, J.M., and Zhong, L. (2009). Duplicated proteasome subunit genes in Drosophila and their roles in spermatogenesis. Heredity 103, 23–31.

10. Betran, E., Thornton, K., and Long, M. (2002). Retroposed new genes out of the X in Drosophila. Genome Res 12, 1854–1859.

11. Brown, J.B., Boley, N., Eisman, R., May, G.E., Stoiber, M.H., Duff, M.O., Booth, B.W., Wen, J.Y., Park, S., Suzuki, A.M., et al. (2014). Diversity and dynamics of the Drosophila transcriptome. Nature 512, 393–399.

12. Buschauer, R., Matsuo, Y., Sugiyama, T., Chen, Y.H., Alhusaini, N., Sweet, T., Ikeuchi, K., Cheng, J.D., Matsuki, Y., Nobuta, R., et al. (2020). The Ccr4-Not complex monitors the translating ribosome for codon optimality. Science 368, 281.

13. Carelli, F.N., Hayakawa, T., Go, Y., Imai, H., Warnefors, M., and Kaessmann, H. (2016). The life history of retrocopies illuminates the evolution of new mammalian genes. Genome Res 26, 301–314.

14. Casola, C., and Betran, E. (2017). The Genomic Impact of Gene Retrocopies: What Have We Learned from Comparative Genomics, Population Genomics, and Transcriptomic Analyses? Genome Biol Evol 9, 1351–1373.

15. Chen, S., Zhang, Y.E., and Long, M. (2010). New genes in Drosophila quickly become essential. Science 330, 1682–1685.

16. Chen, S., Ni, X., Krinsky, B.H., Zhang, Y.E., Vibranovski, M.D., White, K.P., and Long, M. (2012). Reshaping of global gene expression networks and sex-biased gene expression by integration of a young gene. EMBO J 31, 2798–2809.

17. Chen, Y., Boland, A., Kuzuoglu-Ozturk, D., Bawankar, P., Loh, B., Chang, C.T., Weichenrieder, O., and Izaurralde, E. (2014a). A DDX6-CNOT1 Complex and W- Binding Pockets in CNOT9 Reveal Direct Links between miRNA Target Recognition and Silencing. Mol Cell 54, 737–750.

18. Chen, Z.X., Sturgill, D., Qu, J.X., Jiang, H.Y., Park, S., Boley, N., Suzuki, A.M., Fletcher, A.R., Plachetzki, D.C., FitzGerald, P.C., et al. (2014b). Comparative validation of the D. melanogaster modENCODE transcriptome annotation. Genome Res 24, 1209–1223.

19. Clark, A.G., Eisen, M.B., Smith, D.R., Bergman C.M., Oliver, B., et al. (2007). Evolution of genes and genomes on the Drosophila phylogeny. Nature 450, pp. 203–218.

20. Collart, M.A. (2016). The Ccr4-Not complex is a key regulator of eukaryotic gene expression. Wires Rna 7, 438–454.

21. Collart, M.A., and Panasenko, O.O. (2017). The Ccr4-Not Complex: Architecture and Structural Insights. Macromolecular Protein Complexes: Structure and Function 83, 349–379.

22. Dai, H.Z., Yoshimatsu, T.F., and Long, M.Y. (2006). Retrogene movement within- and between-chromosomes in the evolution of Drosophila genomes. Gene 385, 96–102.

23. Ding, Y., Zhao, L., Yang, S., Jiang, Y., Chen, Y., Zhao, R., Zhang, Y., Zhang, G., Dong, Y., Yu, H., et al. (2010). A young Drosophila duplicate gene plays essential roles in spermatogenesis by regulating several Y-linked male fertility genes. Plos Genet 6, e1001255.

24. Dobin, A., Davis, C.A., Schlesinger, F., Drenkow, J., Zaleski, C., Jha, S., Batut, P., Chaisson, M., and Gingeras, T.R. (2013). STAR: ultrafast universal RNA-seq aligner. Bioinformatics 29, 15–21.

25. Eden, E., Navon, R., Steinfeld, I., Lipson, D., and Yakhini, Z. (2009). GOrilla: a tool for discovery and visualization of enriched GO terms in ranked gene lists. BMC Bioinformatics 10, 1–7.

26. Edgar, R.C. (2004). MUSCLE: a multiple sequence alignment method with reduced time and space complexity. BMC Bioinformatics 5, 1–19.

27. Emerson, J.J., Kaessmann, H., Betran, E., and Long, M.Y. (2004). Extensive gene traffic on the mammalian X chromosome. Science 303, 537–540.

28. Erwin, D.H., and Davidson, E.H. (2009). The evolution of hierarchical gene regulatory networks. Nat Rev Genet 10, 141–148.

29. Garapaty, S., Mahajan, M.A., and Samuels, H.H. (2008). Components of the CCR4- NOT complex function as nuclear hormone receptor coactivators via association with the NRC-interacting factor NIF-1. J Biol Chem 283, 6806–6816.

30. Garces, R.G., Gillon, W., and Pai, E.F. (2007). Atomic model of human Rcd-1 reveals an armadillo-like-repeat protein with in vitro nucleic acid binding properties. Protein Sci 16, 176–188.

31. Gnad, F., and Parsch, J. (2006). Sebida: a database for the functional and evolutionary analysis of genes with sex-biased expression. Bioinformatics 22, 2577–2579.

32. Grant, B.J., Rodrigues, A.P., ElSawy, K.M., McCammon, J.A. and Caves, L.S., 2006. Bio3d: an R package for the comparative analysis of protein structures. Bioinformatics, 22(21), pp.2695–2696

33. Halfon, M.S. (2017). Perspectives on Gene Regulatory Network Evolution. Trends Genet 33, 436–447.

34. Harrison, P.W., Wright, A.E., Zimmer, F., Dean, R., Montgomery, S.H., Pointer, M.A., and Mank, J.E. (2015). Sexual selection drives evolution and rapid turnover of male gene expression. Proc Natl Acad Sci 112, 4393–4398.

35. Hiller, M., Chen, X., Pringle, M.J., Suchorolski, M., Sancak, Y., Viswanathan, S., Bolival, B., Lin, T.Y., Marino, S., and Fuller, M.T. (2004). Testis-specific TAF homologs collaborate to control a tissue-specific transcription program. Development 131, 5297–5308.

36. Jiang, L., Li, T., Zhang, X.X., Zhang, B.B., Yu, C.P., Li, Y., Fan, S.X., Jiang, X.H., Khan, T., Hao, Q.M., et al. (2017). RPL10L Is Required for Male Meiotic Division by Compensating for RPL10 during Meiotic Sex Chromosome Inactivation in Mice. Curr Bio. 27, 1498–1505.

37. Kaessmann, H., Vinckenbosch, N., and Long, M. (2009). RNA-based gene duplication: mechanistic and evolutionary insights. Nat Rev Genet 10, 19–31.

38. Kaessmann, H. (2010). Origins, evolution, and phenotypic impact of new genes. Genome Res 20, 1313–1326.

39. Kamath, R.S., Fraser, A.G., Dong, Y., Poulin, G., Durbin, R., Gotta, M., Kanapin, A., Le Bot, N., Moreno, S., Sohrmann, M., et al. (2003). Systematic functional analysis of the Caenorhabditis elegans genome using RNAi. Nature 421, 231–237.

40. Kasinathan, B., Colmenares III, S.U., McConnell, H., Young, J.M., Karpen, G.H. and Malik, H.S., (2020). Innovation of heterochromatin functions drives rapid evolution of essential ZAD-ZNF genes in Drosophila. Elife 9, p.e63368.

41. Keskeny, C., Raisch, T., Sgromo, A., Igreja, C., Bhandari, D., Weichenrieder, O., and Izaurralde, E. (2019). A conserved CAF40-binding motif in metazoan NOT4 mediates association with the CCR4–NOT complex. Genes Dev. 33(3-4), 236– 252.

42. Kondrashov, F.A. (2012). Gene duplication as a mechanism of genomic adaptation to a changing environment. P Roy Soc B-Biol Sci 279, 5048–5057.

43. Kumar, S., Stecher, G., Li, M., Knyaz, C., and Tamura, K. (2018). MEGA X: Molecular Evolutionary Genetics Analysis across Computing Platforms. Mol Biol Evol 35, 1547–1549.

44. Lee, Y.C.G., Ventura, I.M., Rice, G.R., Chen, D.Y., Colmenares, S.U., and Long, M. (2019). Rapid Evolution of Gained Essential Developmental Functions of a Young Gene via Interactions with Other Essential Genes. Mol Biol Evol 36, 2212–2226.

45. Legrand, J.M.D., and Hobbs, R.M. (2018). RNA processing in the male germline: Mechanisms and implications for fertility. Seminars in cell & developmental biology 79, 80–91.

46. Liao, Y., Smyth, G.K., and Shi, W. (2014). featureCounts: an efficient general purpose program for assigning sequence reads to genomic features. Bioinformatics 30, 923–930.

47. Long, M.Y., and Emerson, J.J. (2017). Meiotic Sex Chromosome Inactivation: Compensation by Gene Traffic. Curr Biol. 27, R659–R661.

48. Long, M.Y., VanKuren, N.W., Chen, S.D., and Vibranovski, M.D. (2013). New Gene Evolution: Little Did We Know. Annu Rev Genet 47, 307–333.

49. Love, M.I., Huber, W., and Anders, S. (2014). Moderated estimation of fold change and dispersion for RNA-seq data with DESeq2. Genome Biol 15, 550.

50. Mahadevaraju, S., Fear, J.M., Akeju, M., Galletta, B.J., Pinheiro, M.M., Avelino, C.C., Cabral-de-Mello, D.C., Conlon, K., Dell’Orso, S., Demere, Z., et al. (2021). Dynamic Sex Chromosome Expression in Drosophila Male Germ Cells. Nature Comm 12, 1–16.

51. Markow, T. A., O’Grady, P. M. (2007). Drosophila Biology in the Genomic Age. Genetics 177, 1269–1276.

52. Mathys, H., Basquin, J., Ozgur, S., Czarnocki-Cieciura, M., Bonneau, F., Aartse, A., Dziembowski, A., Nowotny, M., Conti, E., and Filipowicz, W. (2014). Structural and biochemical insights to the role of the CCR4-NOT complex and DDX6 ATPase in microRNA repression. Molecular cell 54, 751–765.

53. Matsuno, M., Compagnon, V., Schoch, G.A., Schmitt, M., Debayle, D., Bassard, J.E., Pollet, B., Hehn, A., Heintz, D., Ullmann, P. and Lapierre, C. (2009). Evolution of a novel phenolic pathway for pollen development. Science 325(5948), pp.1688–1692.

54. Miklos, G.L., and Rubin, G.M. (1996). The role of the genome project in determining gene function: insights from model organisms. Cell 86, 521–529.

55. Miller, J.E., and Reese, J.C. (2012). Ccr4-Not complex: the control freak of eukaryotic cells. Crit Rev Biochem Mol Biol 47, 315–333.

56. Mirny, L.A. and Shakhnovich, E. I. (1999). Universally conserved positions in protein folds: reading evolutionary signals about stability, folding kinetics and function. J Mol Biol 291, 177–196.

57. Ni, J.Q., Zhou, R., Czech, B., Liu, L.P., Holderbaum, L., Yang-Zhou, D., Shim, H.S., Tao, R., Handler, D., Karpowicz, P. and Binari, R. (2011). A genome-scale shRNA resource for transgenic RNAi in Drosophila. Nature methods 8, pp.405–407.

58. Perkins, L.A., Holderbaum, L., Tao, R., Hu, Y., Sopko, R., McCall, K., Yang-Zhou, D., Flockhart, I., Binari, R., Shim, H.-S., et al. (2015). The Transgenic RNAi Project at Harvard Medical School: Resources and Validation. Genetics 201, 843–852.

59. Posada, D. (2008). jModelTest: phylogenetic model averaging. Mol Biol Evol 25, 1253–1256.

60. Quezada-Diaz, J.E., Muliyil, T., Rio, J., and Betran, E. (2010). Drcd-1 related: a positively selected spermatogenesis retrogene in Drosophila. Genetica 138, 925–937.

61. Rehwinkel, J., Behm-Ansmant, I., Gatfield, D., and Izaurralde, E. (2005). A crucial role for GW182 and the DCP1:DCP2 decapping complex in miRNA-mediated gene silencing. RNA 11, 1640–1647.

62. Richler, C., Soreq, H., and Wahrman, J. (1992). X-Inactivation in Mammalian Testis Is Correlated with Inactive X-Specific Transcription. Nat Genet 2, 192–195.

63. Ritchie, M.E., Phipson, B., Wu, D., Hu, Y.F., Law, C.W., Shi, W., and Smyth, G.K. (2015). limma powers differential expression analyses for RNA-sequencing and microarray studies. Nucleic Acids Res 43, e47–e47.

64. Raudvere, U., Kolberg, L., Kuzmin, I., Arak, T., Adler, P., Peterson, H. and Vilo, J., (2019). g: Profiler: a web server for functional enrichment analysis and conversions of gene lists (2019 update). Nucleic Acids Res 47, W191–W198.

65. Robinson, M.D., McCarthy, D.J., and Smyth, G.K. (2010). edgeR: a Bioconductor package for differential expression analysis of digital gene expression data. Bioinformatics 26, 139–140.

66. Ronquist, F., Teslenko, M., van der Mark, P., Ayres, D.L., Darling, A., Hohna, S., Larget, B., Liu, L., Suchard, M.A., and Huelsenbeck, J.P. (2012). MrBayes 3.2: Efficient Bayesian Phylogenetic Inference and Model Choice Across a Large Model Space. Syst Biol 61, 539–542.

67. Ross, B.D., Rosin, L., Thomae, A.W., Hiatt, M.A., Vermaak, D., de la Cruz, A.F., Imhof, A., Mellone, B.G., and Malik, H.S. (2013). Stepwise evolution of essential centromere function in a Drosophila neogene. Science 340, 1211–1214.

68. Russo, C. A., Takezaki, N., & Nei, M. (1995). Molecular phylogeny and divergence times of drosophilid species. Mol Biol Evol 12(3), 391–404.

69. Saleem, S., Schwedes, C.C., Ellis, L.L., Grady, S.T., Adams, R.L., Johnson, N., Whittington, J.R., and Carney, G.E. (2012). Drosophila melanogaster p24 trafficking proteins have vital roles in development and reproduction. Mechanisms of development 129, 177–191.

70. Shannon, C. E. (1948). A Mathematical Theory of Communication. The System Technical J. 27, 379–422.

71. Sgromo, A., Raisch, T., Bawankar, P., Bhandari, D., Chen, Y., Kuzuoglu-Ozturk, D., Weichenrieder, O., and Izaurralde, E. (2017). A CAF40-binding motif facilitates recruitment of the CCR4-NOT complex to mRNAs targeted by Drosophila Roquin. Nat Commun 8, 1–16.

72. Sgromo, A., Raisch, T., Backhaus, C., Keskeny, C., Alva, V., Weichenrieder, O., and Izaurralde, E. (2018). Drosophila Bag-of-marbles directly interacts with the CAF40 subunit of the CCR4-NOT complex to elicit repression of mRNA targets. RNA 24, 381–395.

73. Toups, M.A., and Hahn, M.W. (2010). Retrogenes Reveal the Direction of Sex-Chromosome Evolution in Mosquitoes. Genetics 186, 763–766.

74. Tritschler, F., Eulalio, A., Truffault, V., Hartmann, M.D., Helms, S., Schmidt, S., Coles, M., Izaurralde, E., and Weichenrieder, O. (2007). A divergent Sm fold in EDC3 proteins mediates DCP1 binding and P-body targeting. Mol Cell Biol 27, 8600–8611.

75. Tritschler, F., Eulalio, A., Helms, S., Schmidt, S., Coles, M., Weichenrieder, O., Izaurralde, E., and Truffault, V. (2008). Similar modes of interaction enable Trailer Hitch and EDC3 to associate with DCP1 and Me31B in distinct protein complexes. Mol Cell Biol 28, 6695–6708.

76. VanKuren, N.W., and Vibranovski, M.D. (2014). A novel dataset for identifying sex-biased genes in Drosophila. J Genomics 2, 64–67.

77. VanKuren, N.W., and Long, M.Y. (2018). Gene duplicates resolving sexual conflict rapidly evolved essential gametogenesis functions. Nat Ecol Evol 2, 705–712.

78. Vibranovski, M.D., Lopes, H.F., Karr, T.L., and Long, M. (2009a). Stage-specific expression profiling of Drosophila spermatogenesis suggests that meiotic sex chromosome inactivation drives genomic relocation of testis-expressed genes. Plos Genet 5, e1000731.

79. Vibranovski, M.D., Zhang, Y., and Long, M. (2009b). General gene movement off the X chromosome in the Drosophila genus. Genome Res 19, 897–903.

80. Vibranovski, M.D., Zhang, Y.E., Kemkemer, C., Lopes, H.F., Karr, T.L., and Long, M. (2012). Re-analysis of the larval testis data on meiotic sex chromosome inactivation revealed evidence for tissue-specific gene expression related to the drosophila X chromosome. BMC Biol 10, 49.

81. Mering, C.V., Huynen, M., Jaeggi, D., Schmidt, S., Bork, P. and Snel, B. (2003). STRING: a database of predicted functional associations between proteins. Nucleic Acids Res 31(1), pp.258–261.

82. Wahle, E., and Winkler, G. S., (2013). RNA decay machines: deadenylation by the CCR4–NOT and Pan2-Pan3 complexes. Biochim Bio-phys Acta 1829, 561–570.

83. Wang, T., Birsoy, K., Hughes, N.W., Krupczak, K.M., Post, Y., Wei, J.J., Lander, E.S., and Sabatini, D.M. (2015). Identification and characterization of essential genes in the human genome. Science 350, 1096–1101.

84. White-Cooper, H. (2010). Molecular mechanisms of gene regulation during Drosophila spermatogenesis. Reproduction 139, 11–21.

85. White-Cooper, H. (2012). Tissue, cell type and stage-specific ectopic gene expression and RNAi induction in the Drosophila testis. Spermatogenesis 2, 11–22.

86. Witt, E., Benjamin, S., Svetec, N., and Zhao, L. (2019). Testis single-cell RNA-seq reveals the dynamics of de novo gene transcription and germline mutational bias in Drosophila. Elife 8, e47138.

87. Wu, C.I., and Xu, E.Y., (2003). Sexual antagonism and X inactivation–the SAXI hypothesis. TRENDS in Genetics 19, 243–247.

88. Xia, S., VanKuren, N.W., Chen, C., Zhang, L., Kemkemer, C., Shao, Y., Jia, H., Lee, U., Advani, A.S., Gschwend, A. and Vibranovski, M.D., (2021). Genomic analyses of new genes and their phenotypic effects reveal rapid evolution of essential functions in Drosophila development. PLoS genetics 17, p.e1009654.

89. Zekri, L., Kuzuoglu-Ozturk, D., and Izaurralde, E. (2013). GW182 proteins cause PABP dissociation from silenced miRNA targets in the absence of deadenylation. EMBO J 32, 1052–1065.

90. Zhang, W.Y., Landback, P., Gschwend, A.R., Shen, B.R., and Long, M.Y. (2015). New genes drive the evolution of gene interaction networks in the human and mouse genomes. Genome Biol 16, 202.

91. Zhang, Y.E., Vibranovski, M.D., Krinsky, B.H., and Long, M.Y. (2010). Age-dependent chromosomal distribution of male-biased genes in Drosophila. Genome Res 20, 1526–1533.

92. Zhong, L., and Belote, J.M. (2007). The testis-specific proteasome subunit Pros alpha 6T of D-melanogaster is required for individualization and nuclear maturation during spermatogenesis. Development 134, 3517–3525.

