## Supplementary figure for "Rapid gene evolution in an ancient post-transcriptional and translational regulatory system compensates for meiotic X chromosomal inactivation"

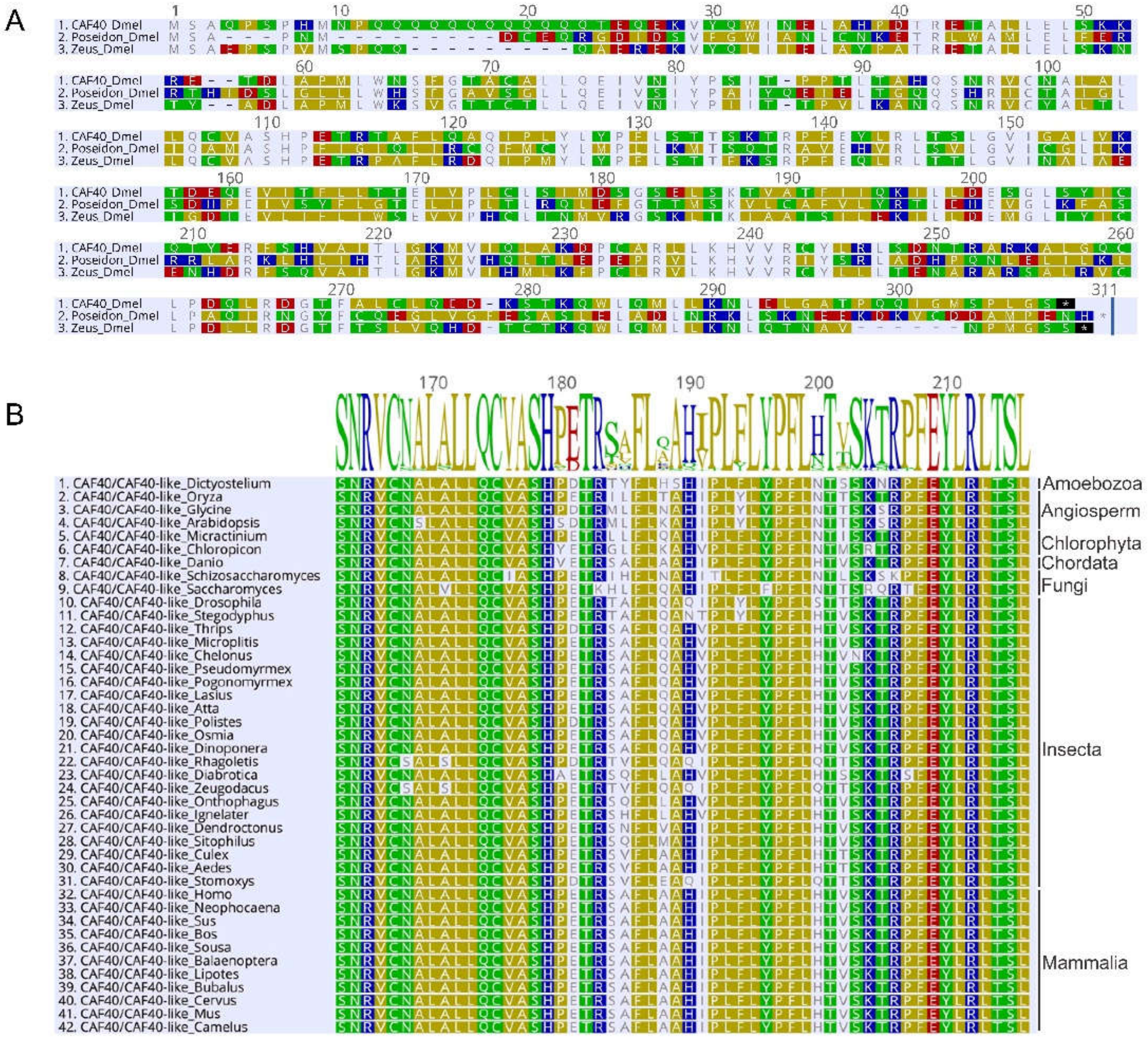
A
1. CAF40_Dmel
2. Poseidon_Dmel
3. Zeus_Dmel
B
SNRVCNALALLQCVASHPETRSAFLQAHIPLFLYPFLHTVSKTRPFEYLRLTSL
1. CAF40/CAF40-like_Dictyostelium
2. CAF40/CAF40-like_Oryza
3. CAF40/CAF40-like_Glycine
4. CAF40/CAF40-like_Arabidopsis
5. CAF40/CAF40-like_Micractinium
6. CAF40/CAF40-like_Chloropicon
7. CAF40/CAF40-like_Danio
8. CAF40/CAF40-like_Schizosaccharomyces
9. CAF40/CAF40-like_Saccharomyces
10. CAF40/CAF40-like_Drosophila
11. CAF40/CAF40-like_Stegodyphus
12. CAF40/CAF40-like_Thrips
13. CAF40/CAF40-like_Microplitis
14. CAF40/CAF40-like_Chelonus
15. CAF40/CAF40-like_Pseudomyrmex
16. CAF40/CAF40-like_Pogonomyrmex
17. CAF40/CAF40-like_Lasius
18. CAF40/CAF40-like_Atta
19. CAF40/CAF40-like_Polistes
20. CAF40/CAF40-like_Osmia
21. CAF40/CAF40-like_Dinoponera
22. CAF40/CAF40-like_Rhagoletis
23. CAF40/CAF40-like_Diabrotica
24. CAF40/CAF40-like_Zeugodacus
25. CAF40/CAF40-like_Onthophagus
26. CAF40/CAF40-like_Ignelater
27. CAF40/CAF40-like_Dendroctonus
28. CAF40/CAF40-like_Sitophilus
29. CAF40/CAF40-like_Culex
30. CAF40/CAF40-like_Aedes
31. CAF40/CAF40-like_Stomoxys
32. CAF40/CAF40-like_Homo
33. CAF40/CAF40-like_Neophocaena
34. CAF40/CAF40-like_Sus
35. CAF40/CAF40-like_Bos
36. CAF40/CAF40-like_Sousa
37. CAF40/CAF40-like_Balaenoptera
38. CAF40/CAF40-like_Lipotes
39. CAF40/CAF40-like_Bubalus
40. CAF40/CAF40-like_Cervus
41. CAF40/CAF40-like_Mus
42. CAF40/CAF40-like_Camelus
Amoebozoa
Angiosperm
Chlorophyta
Chordata
Fungi
Insecta
Mammalia

C

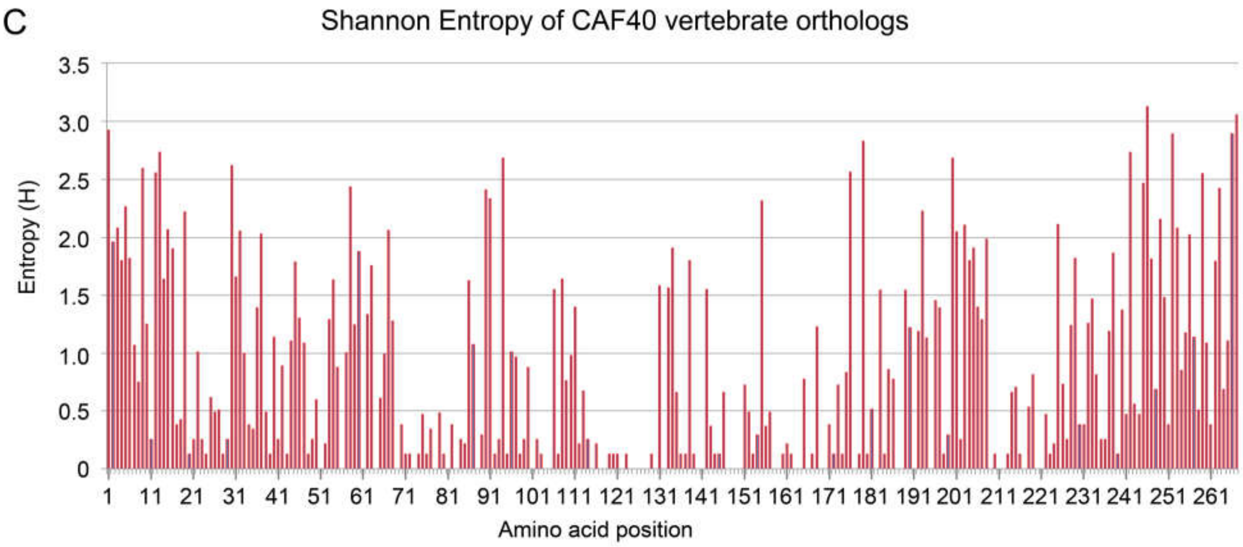


D

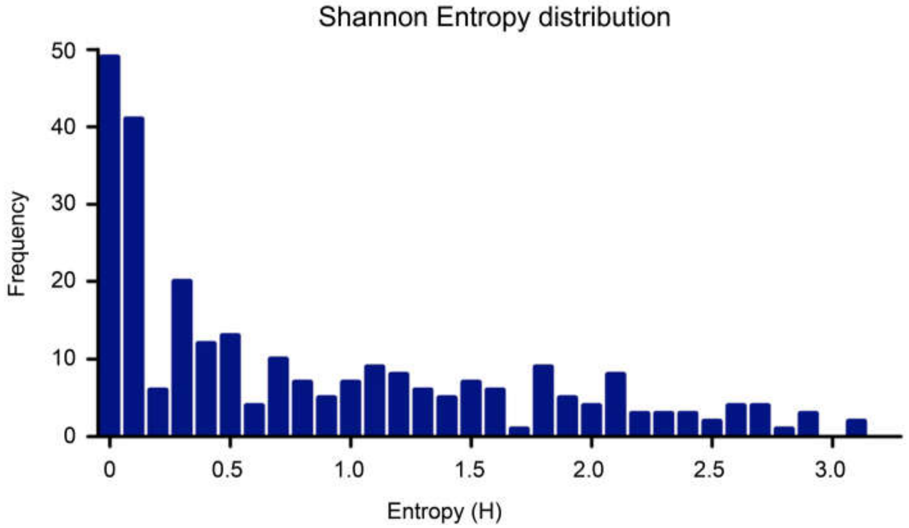


**Figure S1. The divergence and conservation of protein sequences of CAF40, Poseidon and Zeus.**

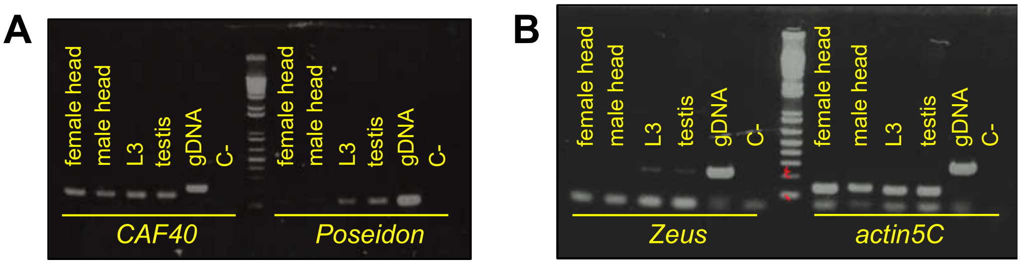


**Figure S2. Agarose gels confirming the expression of each paralog at different tissues from *D. melanogaster*.**

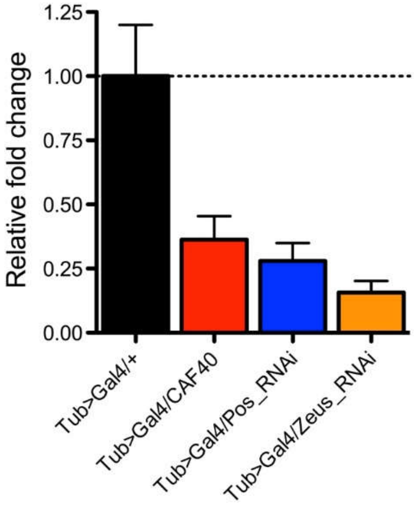


**Figure S3. Constitutive RNAi-knockdown efficiency measured by quantitative PCR.**

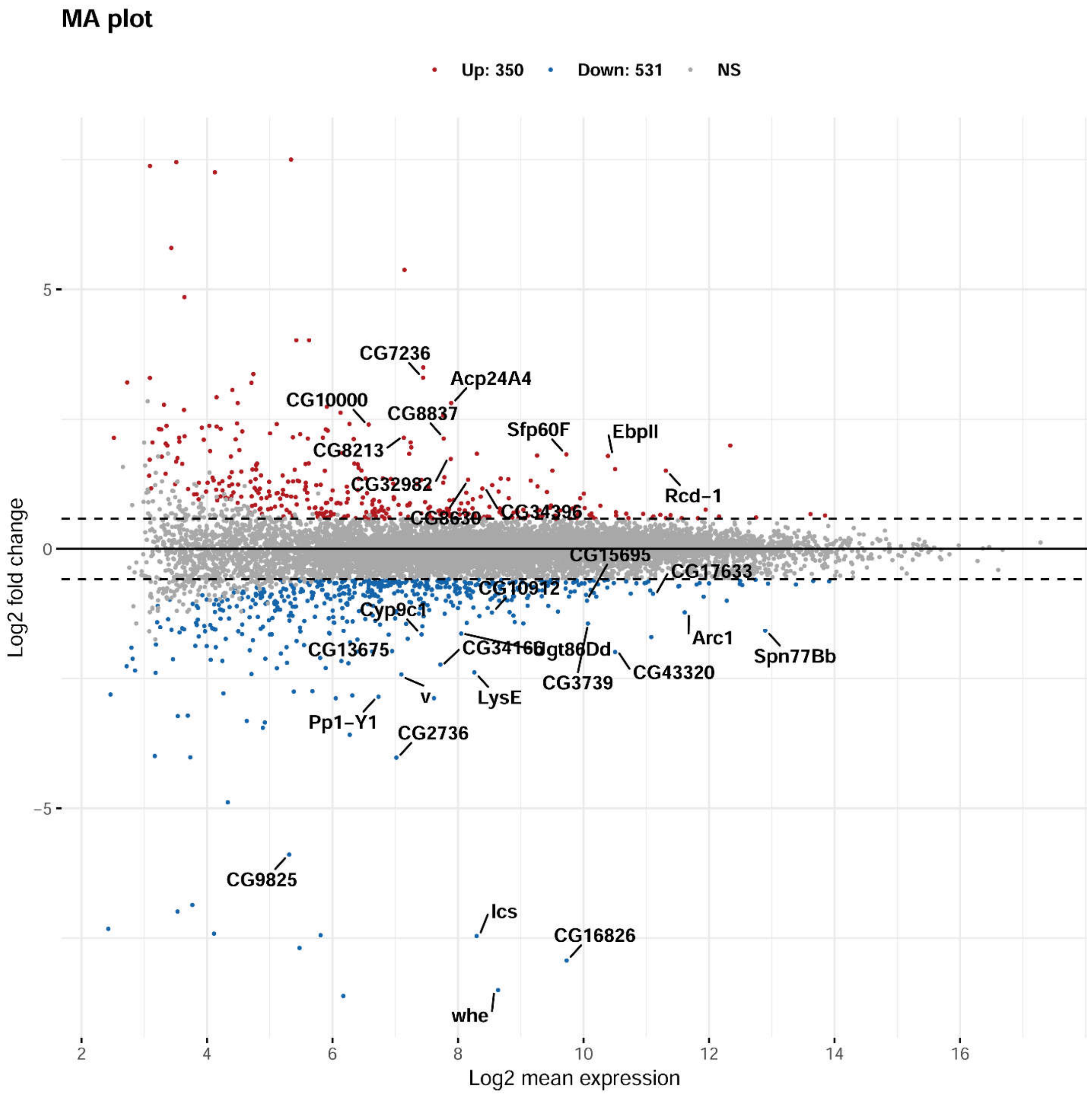


**Figure S4. MA plot of DEGs between Poseidon-KD and CAF40-KD line by deseq2.**

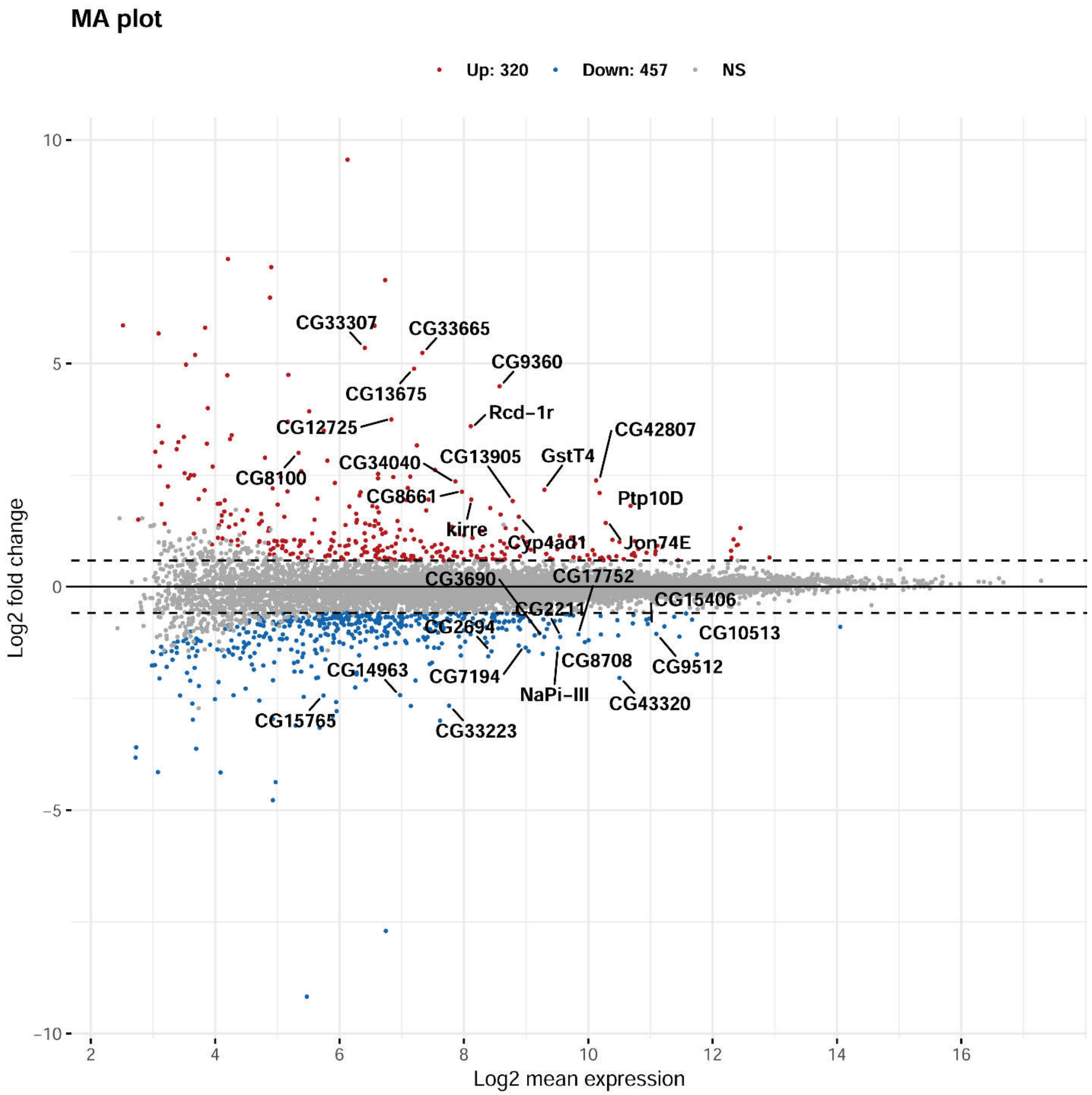


**Figure S5. MA plot of DEGs between Poseidon-KD and Zeus-KD line by deseq2.**

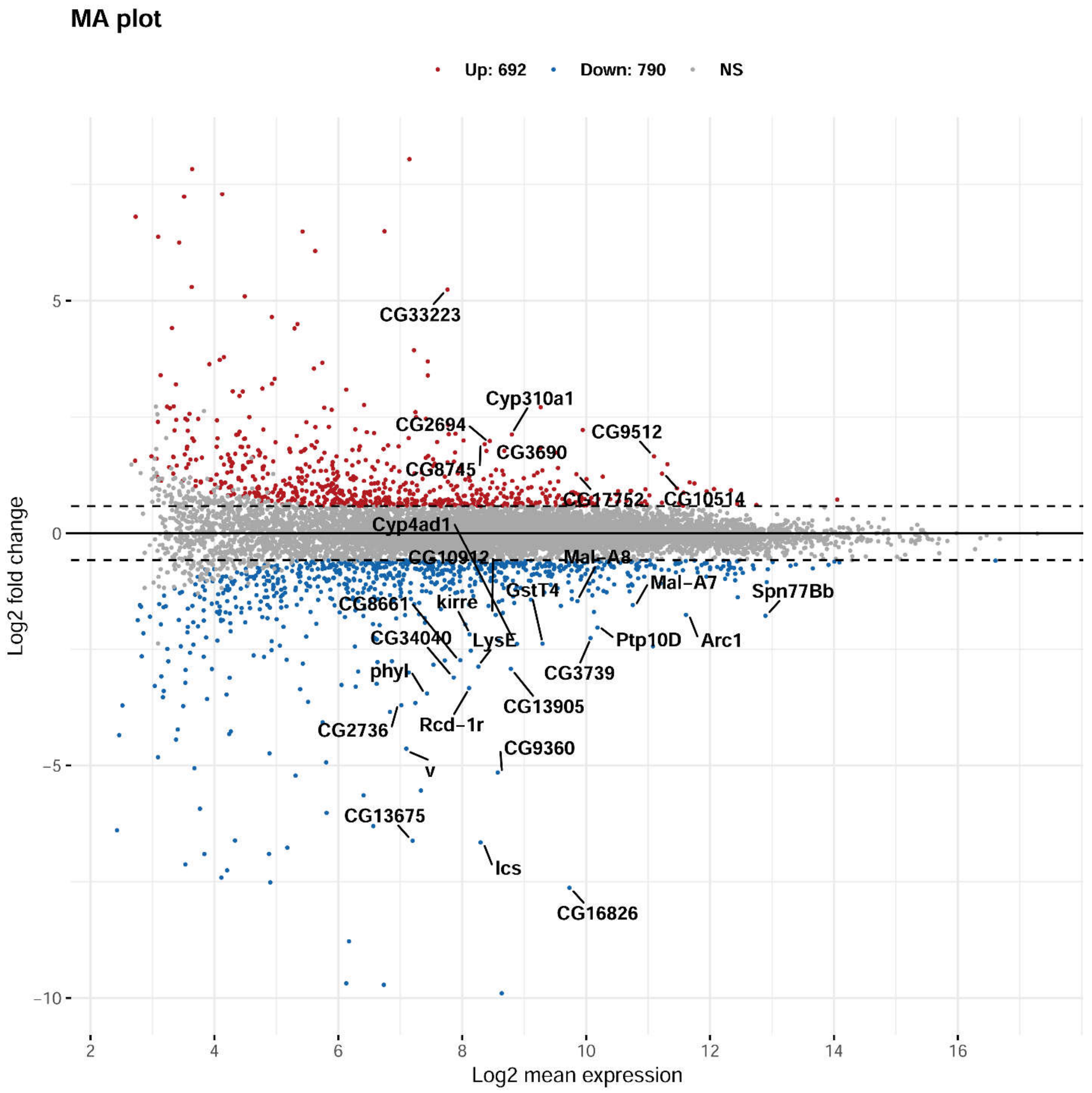


**Figure S6. MA plot of DEGs between Zeus-KD and CAF40-KD line by deseq2.**

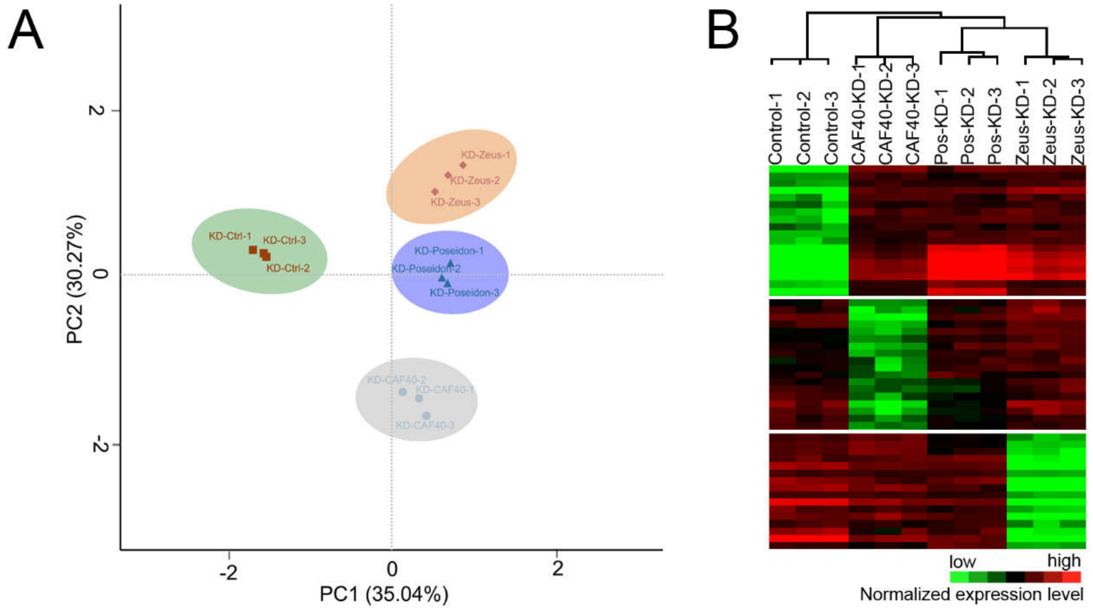


**Figure S7. Clustering of samples upon knockdown in the RNA-seq assay.**

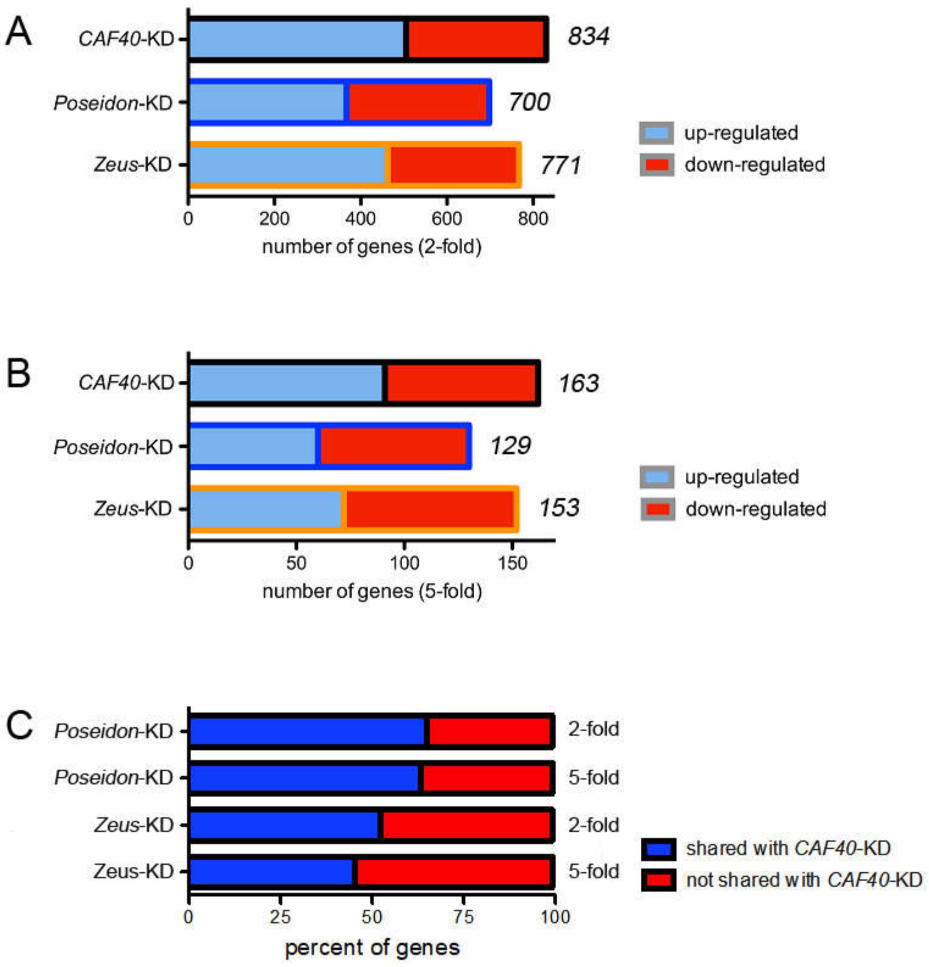


**Figure S8. Impact of *CAF40*, *Poseidon* and *Zeus* knockdowns on global gene expression (A. and B.) and Proportion of genes differentially expressed upon *Poseidon* and *Zeus* knockdown that are also impacted by *CAF40* (C.).**

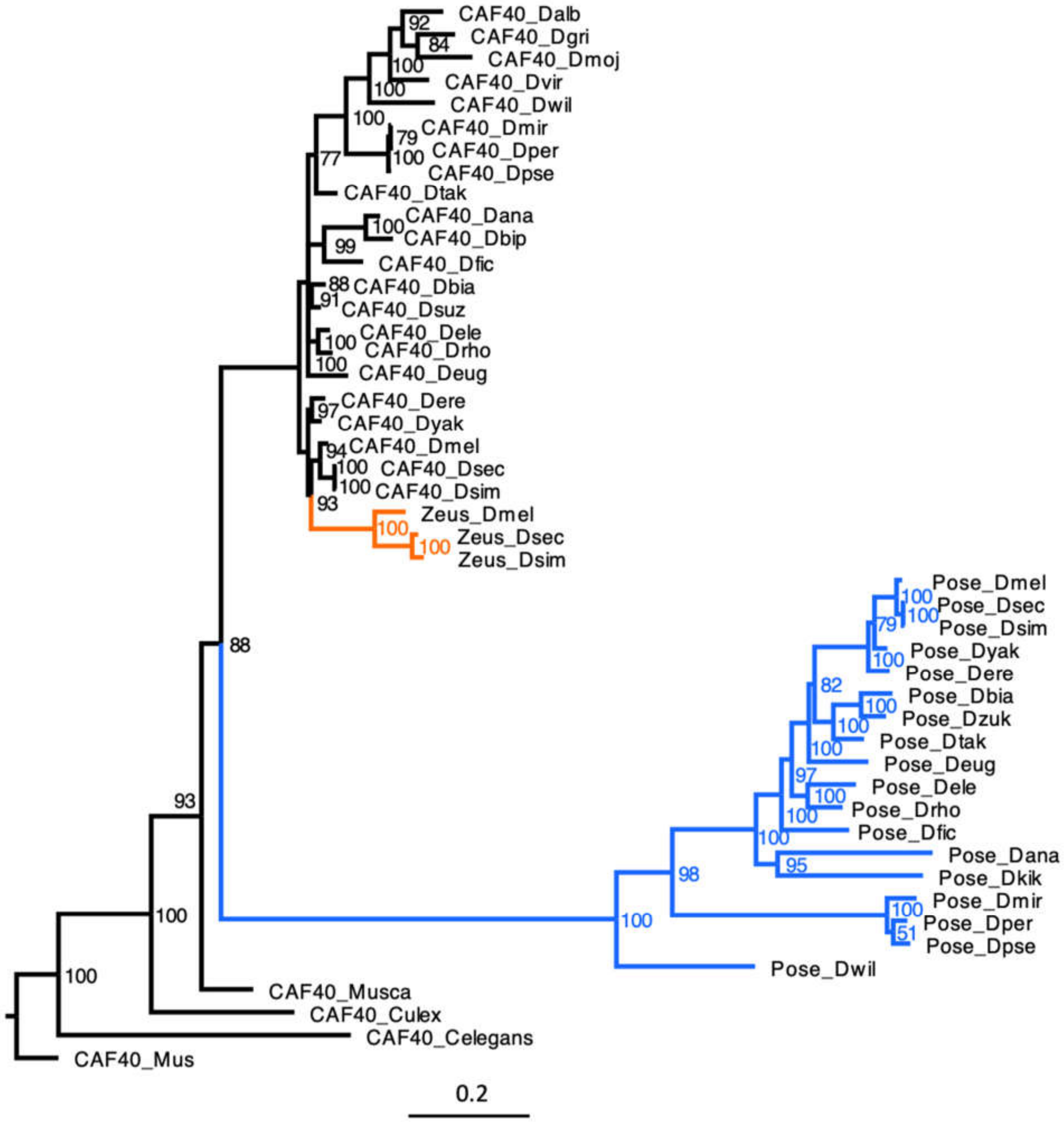


**Figure S9. Phylogenetic relationship among the three paralogs reconstructed through Bayesian method. *CAF40* homologs in black, *Poseidon* in blue, and *Zeus* in orange; numbers represent the Bayesian posterior probability of each node.**

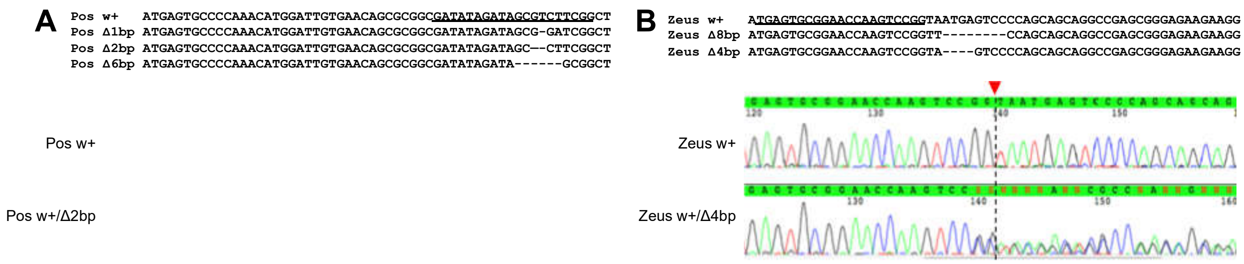


**Figure S10. Knockout mutants for *Poseidon* (A) and *Zeus* (B) generated through CRISPR-Cas9.**

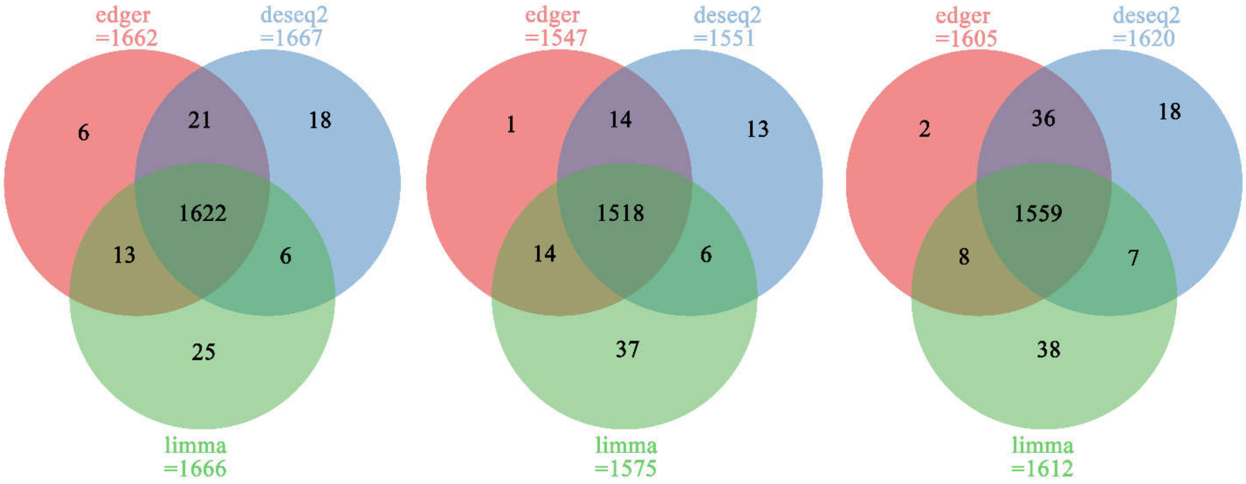


**Figure S11. DEGs analysis of CAF40-KD vs Control, Poseidon-KD vs Control and Zeus-KD vs Control by using edger, deseq2, and limma.**
